# Cryo-EM reveals multiple mechanisms of ribosome inhibition by doxycycline

**DOI:** 10.1101/2025.11.08.686982

**Authors:** William S. Stuart, Michail N. Isupov, Mathew McLaren, Christopher H. Jenkins, Adam Monier, Bertram Daum, Isobel H. Norville, Vicki A.M. Gold, Nicholas J. Harmer

## Abstract

Antimicrobial resistance is driving the search for new antibiotics and a greater understanding of their mechanism of action. Doxycycline is amongst the most-prescribed antimicrobials. It demonstrates a particularly low minimum inhibitory concentration against the zoonotic pathogen *Coxiella burnetii*. Doxycycline canonically targets the bacterial ribosome by blocking tRNA binding at the decoding centre (A-site) of the small subunit. Using cryo electron-microscopy, we analysed doxycycline binding to *C. burnetii* and *Escherichia coli* ribosomes. Both structures reveal unexpected binding at the exit tunnel in the large subunit. In *C. burnetii* three doxycycline molecules stack to block the tunnel. In *E. coli* one doxycycline molecule triggers a major uncharacterised conformation of the ribosome. This fundamentally reorganises the peptidyl transferase centre and blocks tRNA binding, challenging the concept that this region is largely static. We identify a new ribosomal protein in the *C. burnetii* large subunit and characterise an additional member of the prokaryotic ribosome hibernation promoting factor family. These insights into ribosome function and antibiotic action may aid the development of new ribosome inhibitor antibiotics.

## Main

The obligate pathogen *Coxiella burnetii* infects a wide range of mammals and other species [1, 2]. With an infectious dose of approximately one bacterium, *C. burnetii* is highly contagious, putting veterinary and farm workers at risk [3–5]. *C. burnetii* has a biphasic lifecycle with large and small cell variants (LCV and SCV, respectively) [6, 7]. The LCV is metabolically active, whereas the quiescent SCV has a spore-like morphology, enabling airborne spread [8]. Over 4,000 human cases were identified during a three year outbreak in the Netherlands (2007 to 2010) [9], with associated costs estimated at up to 600 million Euros [10]. Q-fever, the symptomatic *C. burnetii* infection in humans, initially presents with fever-like symptoms, with up to 5% of symptomatic infections developing into long-term chronic illness, although it can be many years before chronic infections are identified [11, 12]. *C. burnetii* has been named as one of the 24 pathogens posing the greatest risk to public health [13].

Clearance of *C. burnetii* is highly challenging due to its slow growth rate and unique intracellular location within the *Coxiella* containing vacuole (CCV) [14]. Doxycycline (Fig. 1) and hydroxychloroquine are given in a regimen lasting for at least 18 months, consisting of two doses of doxycycline and three doses of hydroxychloroquine (respectively 200 and 300 mg daily) [15, 16]. Hydroxychloroquine likely alkalises the acidic CCV, improving doxycycline performance [17]. *C. burnetii* is exquisitely sensitive to doxycycline (minimum inhibitory concentration (MIC) of between 0.01-0.04 µg/ml and a bactericidal concentration of 8 µg/ml) [18]. In comparison, the MIC of doxycycline against *E. coli* is around 1.5 µg/ml [19].

**Figure 1:**
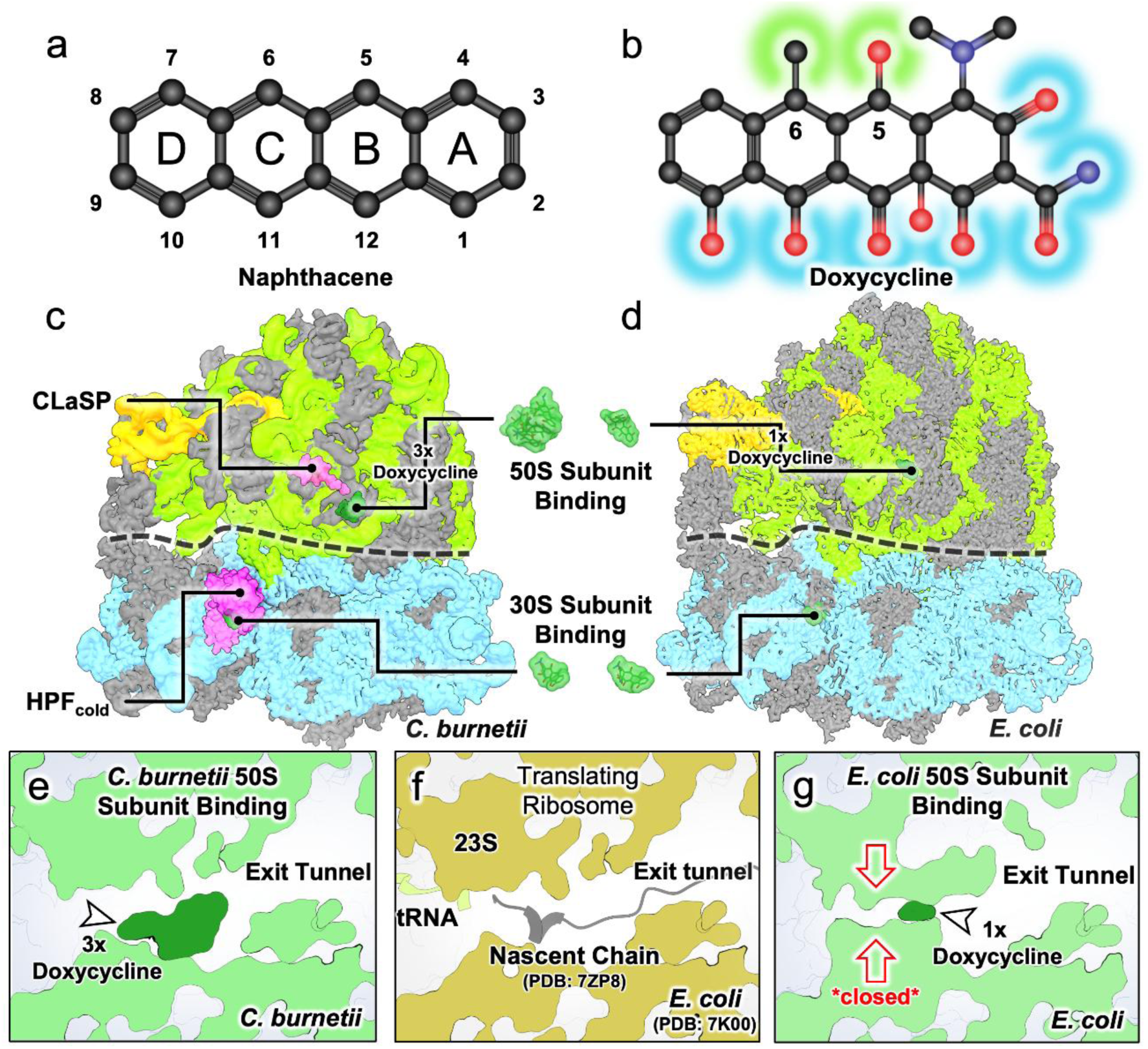
Cryo-EM reveals multiple doxycycline binding sites on the prokaryotic ribosome, a ribosome hibernation factor and a *Coxiellacae* specific ribosomal protein. (**a**) Tetracyclines share a naphthacene core consisting of four fused carbon rings. (**b**) This is decorated by common modifications on carbons 1-3 and 10-12 (highlighted blue). Unique modifications at other positions (typically, carbons 4-8), such as a C5 hydroxyl group and a C6 methyl group for doxycycline (highlighted green), define the different family members. Later generation synthetic tetracyclines focused on additions around the D-ring (C7-9), expanding the physical footprint of the drugs to overcome tetracycline resistance mechanisms [89]. (**c,d**) Overview of the structures of the *Coxiella burnetii* (**c**) and *Escherichia coli* (**d**) ribosomes in complex with doxycycline. Proteins are coloured grey, 16S and 23S rRNA are coloured blue and green respectively and 5S is coloured pale orange. Bound doxycycline is indicated and coloured dark green; bound HPFcold is shown in dark blue (lower left); CLaSP (*Coxiellacae* Large Subunit Peptide) is shown in magenta. (**e,f,g**) Doxycycline blocks the translating ribosome, each shows a cross section through the nascent chain exit tunnel region. The *C. burnetii* ribosome (**e**) is blocked by three molecules of doxycycline that block the tunnel that the nascent chain exits through. For comparison, an active ribosome (**f**), showing the canonical structure (dark yellow, pdb_00007k00, [73]) and nascent chain (grey cartoon, pdb_00007zp8, [90]) in an unimpeded exit tunnel. Doxycycline remodels the r23S RNA chain at the peptidyl transfer centre (**g**), constricting this location and the exit tunnel.

We present the structure of the *C. burnetii* ribosome and compare the doxycycline-bound forms of the *C. burnetii* and *E. coli* ribosomes to identify conformational differences associated with antibiotic binding. Ribosomes have three binding sites for tRNA molecules: the A, P and E-sites. mRNA aligns with the small subunit of the ribosome at the Shine-Dalgarno sequence and enters through the mRNA entrance tunnel. At the A-site the tRNA anti-codon pairs to the mRNA. Amino acids are moved from the tRNA to the nascent chain at the P-site and the extending polypeptide exits through the nascent chain exit tunnel in the large subunit. tRNAs move to the E-site and leave the ribosome. The canonical tetracycline binding site overlaps the A-site tRNA, and we observe doxycycline bound here in each ribosome. However, in both organisms, unexpected doxycycline binding sites were identified around the nascent chain exit tunnel. Exit tunnel binding differs between both species and explains *C. burnetii’s* sensitivity to doxycycline. In the non-doxycycline inhibited forms, paused, hibernating ribosomes predominated, induced by bound hibernation factor proteins. Hibernation factors, found in all forms of life, provide a way of pausing and protecting ribosomes when not engaged in protein synthesis [20, 21]. Since cells eventually transition back to the SCV state within the infected host, where hibernating ribosomes likely predominate, this form is of clinical interest [22, 23]. We observe a previously unknown *C. burnetii* large subunit peptide and identify a subpopulation of *C. burnetii* ribosomes complexed with a protein containing a cold shock domain, which we propose as an additional member of the hibernation promoting factor (HPF) family, named HPF_cold_.

### Doxycycline binds to both large and small ribosomal subunits

To characterise species specific features and contrast antibiotic behaviour, we determined the doxycycline (Fig. 1a-b) bound cryo-EM structures of the *C. burnetii* and *E. coli* ribosomes. Focused classification was used to fully elucidate antibiotic behaviour. Both datasets showed doxycycline bound in the expected 30S subunit location common to other tetracyclines [24–26] (Fig. 1c-d, Fig. S1). Surprisingly, both populations also showed doxycycline bound to the 50S subunit, with distinct modes of binding for the two Gram-negative bacteria (Fig. 1c-g).

### The *Coxiella burnetii* hibernating ribosome

The overall structure of the *C. burnetii* ribosome is similar to other prokaryotic ribosomes (Fig. 1c, Movie 1). Forty-eight proteins identified in the 70S ribosome are listed in Supplementary Table 1 (28 in the large subunit, 20 in the small subunit, including a newly identified large subunit peptide and a small subunit hibernation factor, both discussed below). Any remaining elements of the *C. burnetii* intervening sequence (IVS), found within the 23S ribosomal RNA (rRNA) were not present in the ribosome structure, nor the hypothetical S23 protein (encoded within the precursor r23S rRNA) [27], correlating with previous studies [28]. There are notably fewer rRNA post-translational protein modifications (PTMs) identified for the *C. burnetii* ribosome compared to *E. coli*, with those that remain being mostly in the small subunit (Fig. S2).

### A prokaryotic hibernation factor protects the monomeric ribosome

The *C. burnetii* 70S structure revealed a protein bound to the 30S tRNA cleft (Fig. 2a-b, Movie 1). This was identified as *CBU_0020*, one of two hibernation factors present in the *C. burnetii* genome. *CBU_0020* is a ∼20 kDa protein consisting of two domains joined by a six-residue linker. The N-terminal domain has a sigma-54 fold (Pfam PF02482) and the C-terminal domain a cold shock domain fold (CSD, Pfam PF00313) (Fig. 2b). *CBU_0020* will subsequently be referred to as HPF_cold_.

**Figure 2:**
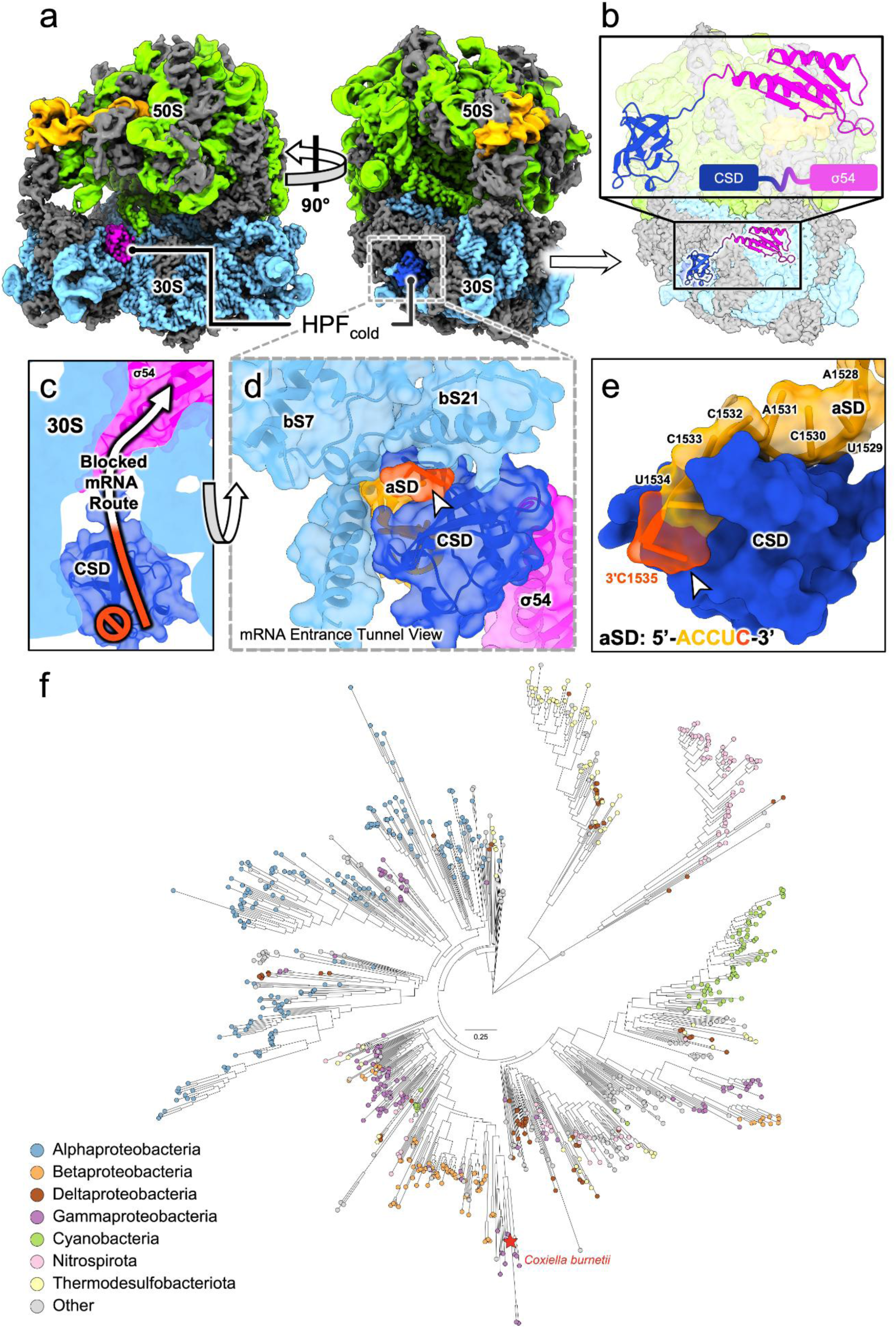
HPF_cold_, a new member of the hibernation factor family. (**a**) A hibernation promoting factor (HPF_cold_; magenta and dark blue) was observed bound within the 30S decoding centre of the *C. burnetii* ribosome. Ribosomal proteins are coloured grey, whilst 5S, 16S and 23S rRNA are coloured light orange, blue and green respectively. (**b**) HPF_cold_ (*CBU_0020*) consists of two domains, an N-terminal sigma-54 modulation factor domain (PF02482, σ54; magenta) and a C-terminal cold-shock domain (PF00313, CSD; dark blue). (**c**) HPF_cold_ occupies the mRNA route through the ribosome, preventing translation. (**d**) The cold shock domain binds the anti-Shine Dalgarno sequence on the 16S rRNA (orange/red). Ribosomal proteins S7 and S21 (pale blue) coordinate the CSD. Proteins shown with semi-transparent surface and cartoon representation. (**e**) The anti-Shine-Dalgarno (aSD) sequence (ACCUC) makes extensive interactions with the CSD. The aSD is at the 3’ end of the r16S and so is susceptible to RNAse activity if not protected. CSD shown as surface, r16S rRNA as semi-transparent surface and cartoon representation. (**f**) Maximum-likelihood phylogeny of fused PF02482–PF00313 two-domain proteins across bacterial lineages. Unrooted ML tree of proteins containing both Pfam domains PF02482 (σ54) and PF00313 (CSD). Tips are coloured by major lineages (Alphaproteobacteria, Betaproteobacteria, Gammaproteobacteria, Deltaproteobacteria, Cyanobacteria, Nitrospirota, Thermodesulfobacteriota, Other). *Coxiella* sequences are highlighted. The alignment was built with MAFFT (G-INS-i) [84] and trimmed with trimAl (gappyout) [85]; the best-fit model was selected with ModelFinder [87], and branch support was estimated with 10,000 ultrafast bootstrap replicates in IQ-TREE [86]. The scale bar denotes 0.25 substitutions/site. Tree visualisation was done in iTOL [91].

The HPF_cold_ CSD domain occupies the mRNA entrance tunnel (Fig. 2c), binding and protecting the r16S anti-Shine Dalgarno (aSD) sequence (Fig. 2d-e, S3a-b). The aSD wraps around the CSD making a combination of stacking interactions, hydrogen bonds and charge interactions (Fig. S3b-c). The CSD is also pincered by the α-helical arms of bS21 and contacted by bS7 (Fig. 2d-e, Fig. S3d).

We have identified HPF_cold_ in all major proteobacterial classes, including Cyanobacteria, and several non-proteobacterial phyla (Fig. 2f, Supplementary Discussion). A Protein Data Bank (PDB) BLAST [29, 30] search of the HPF_cold_ CSD revealed no significant sequence similarity to any existing CSD structure. Strong structural similarity to bacterial cold shock proteins is retained, evidenced with a structural alignment RMSD of 1.8 Å against the full *E. coli* CspA, despite low sequence similarity overall (Fig. S4a,d, S5a) [31]. A Foldseek search [32] of *E. coli* CspA against the *C. burnetii* genome returned HPF_cold_ with the highest E-value and no dedicated cold shock protein family members, indicating these have been lost.

### A *C. burnetii* specific ribosome binding protein

During model building, we observed additional density that could not be attributed to any known ribosomal protein in the PDB. This density located to the cleft formed by domains II and V of the r23S rRNA, together with the D-loop of r5S rRNA (Fig. 3a,b; Fig. S6a,b, Movie 1) and was of sufficient quality to assign a 16 amino acid peptide sequence *de-novo* from the cryo-EM map. A BLAST search [30] of this peptide identified a unique hit within the *C. burnetii* genome located one base 3’ downstream from the ribosomal RNA transcription regulator *nusB* (Fig. S6c). This open reading frame encodes a 22-residue peptide (residues 2-17 are visible in the cryo-EM map) unique to *Coxiellaceae*, which we therefore name CLaSP (*<u>C</u>oxiellaceae* <u>La</u>rge <u>S</u>ubunit <u>P</u>eptide). CLaSP appears to stabilise the ribosome by inserting between the r23S and r5S, anchoring the two (Fig. 3c-d).

**Figure 3:**
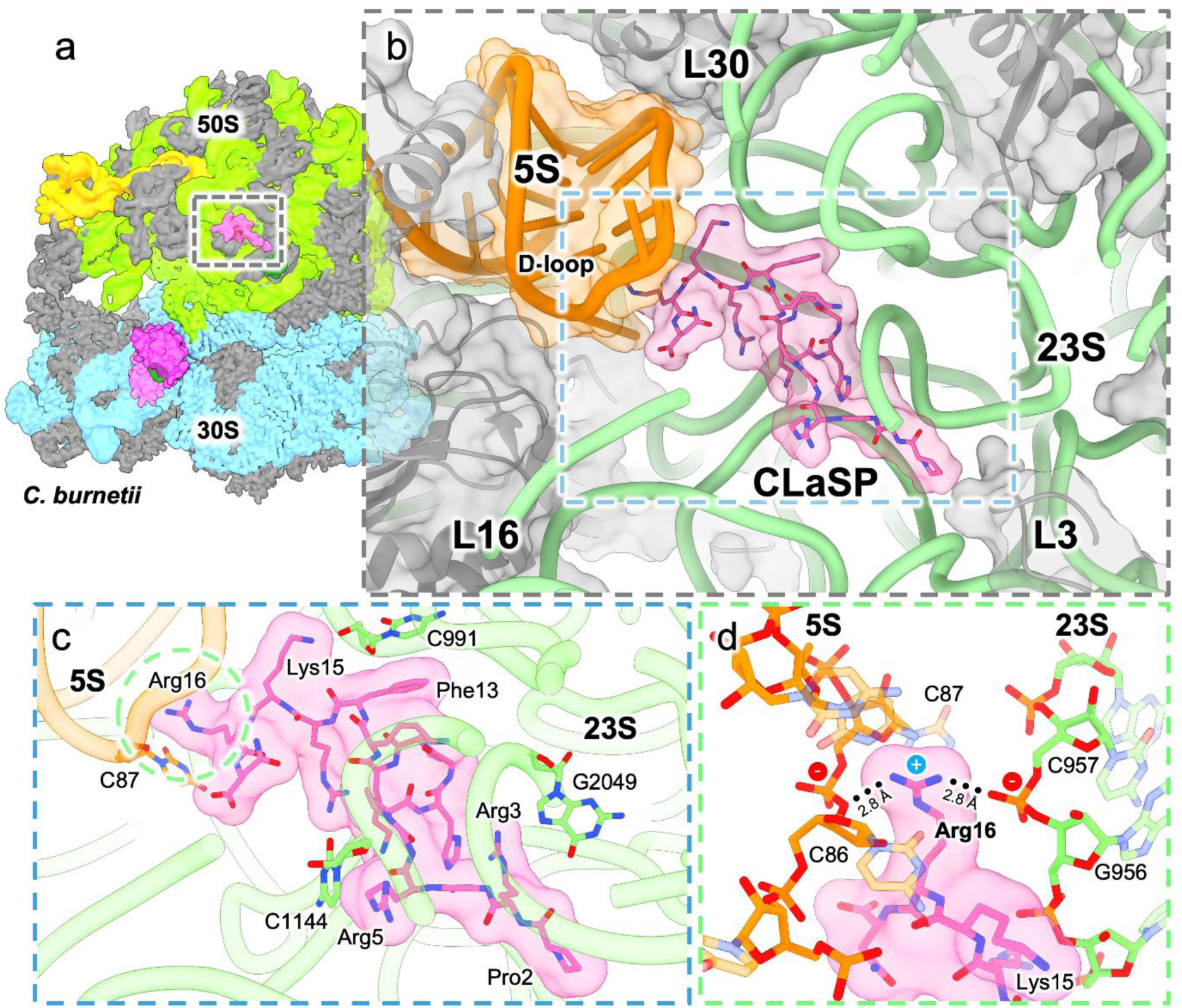
CLaSP: A ribosomal protein unique to *Coxiellacae*. (**a**) Unidentified density in the large subunit was modelled *de-novo* and identified as a new gene, termed CLaSP (*Coxiellaceae* large subunit peptide). Proteins are coloured grey, whilst r16S, r23S and r5S rRNA are coloured blue, green and orange respectively. CLaSP is shown in magenta. (**b**) CLaSP shows an extended conformation and occupies a cleft between the truncated r5S rRNA D-loop and the 23S rRNA. 5S rRNA and ribosomal proteins shown as a transparent surface and cartoon representation in orange and grey respectively, CLaSP as sticks with a transparent surface in pink, and r23S rRNA as green tube. Given the proximity to the PTC, CLaSP likely acts to stabilise the ribosome and may aid in communication within the large subunit. (**c**) CLaSP is a basic peptide, making extensive charge-charge interactions to the surrounding phosphate backbone. Specific stacking interactions are shown with rRNA bases as sticks. (**d**) Arg16 acts as a key coordinating residue, making interactions with the r5S and r23S rRNA backbones. rRNA shown as sticks.

### Doxycycline binds the canonical small subunit site in both *E. coli* and *C. burnetii*

As expected, we observe doxycycline bound in the classical 30S binding site in the tRNA decoding region for both *C. burnetii* and *E. coli* ribosomes (Fig. S1a-b). This binding site is identical to previous 30S tetracycline bound structures, where two coordinating magnesium ions and ring D of doxycycline stacks upon r16S C1054 [24]. This prevents tRNA A-site binding, a mechanism shared with ribosomal hibernation promoting factors. Indeed, in our models, doxycycline overlapped with bound hibernation factors in both species: HPF_cold_ in *C. burnetii* and RaiA in the *E. coli* ribosome. We therefore separated antibiotic bound particles from hibernation factor bound particles for both species using focused classification methods. This improved the density for the bound antibiotic (Fig. S7 and S8) and agrees with the observation in *E. coli* that 30S bound tetracyclines compete directly with tRNA binding site hibernation factors [20].

### Doxycycline blocks the *C. burnetii* nascent peptide tunnel

We observed a remarkable stack of three doxycycline molecules bound in the exit tunnel of the *C. burnetii* ribosome, which we number by increasing distance from the peptidyl transferase centre (PTC; Fig. 4a-c, Fig. S9a, Movie 1). In contrast only incomplete binding of one molecule was observed in the *E. coli* ribosome (Fig S1d). The three-molecule *C. burnetii* binding is an elaboration of a similar 50S binding mode first observed for a single sarecycline molecule against *C. acnes* [33] (Fig. S10c-e). Doxycycline molecule one (DOX1) stacks upon the U1800:U2606 base pair (1782:2586, *E. coli* numbering in parentheses; Fig. S9a). The two other doxycycline molecules stack consecutively on top of the first, with their ‘C’ rings stacked upon the ‘D’ ring of the preceding molecule (Fig. 4c). DOX2 is rotated by approximately 90° around the stack axis with respect to DOX1 and DOX3. This arrangement facilitates interactions with three magnesium ions that are key to the three-molecule binding (Fig. S10a). The central magnesium ion is coordinated by two oxygen atoms from each of the three doxycycline molecules. DOX1 coordinates a second magnesium ion together with U2629 (U2609), while DOX2 and DOX3 coordinate a third magnesium (Fig. 4c, Fig. S10a). The amide group of DOX1 holds in position the flexible ribosome stall sensor U2082 (U2062) in the tunnel protruding conformation (Fig. 4c, Fig. S9e,10a). DOX3 is also closely associated with the edge of the tunnel (Fig. 4c). Here, a magnesium coordinated water forms a H-bond with A2078 (A2058; Fig. S10g). Comparison between bound and unbound ribosomes (Fig. S9a-b) revealed that DOX2 stabilises U2526 (U2506; Fig. S9c-d) and G2077-A2079 (2057-2059) have additional density in the empty state, likely corresponding to bound waters that are displaced on doxycycline binding (Fig. S9f-g). The interaction between DOX2 and U2526 stabilises the base in a luminal, translationally stalled-like position (Fig. S9c). The development of later generation tetracyclines has elaborated the underlying four-ring structure with various chemical decorations (Fig. 4d). We reasoned that the lack of additional chemical elaboration on the doxycycline rings, compared to the bulkier later generation tetracyclines, may facilitate this stacked binding mode (Fig. 4d, Fig. S10b). To test this hypothesis, minimum inhibitory concentration (MIC) experiments were carried out against *C. burnetii* (strain NMII) with four tetracycline derivatives: tetracycline, doxycycline, tigecycline and eravacycline. Tigecycline and eravacycline are third and fourth generation tetracyclines respectively, with chemical additions on C9 (Fig. 4d, Fig. S10c-f). Doxycycline retained the lowest MIC at 0.016 µg/ml and together with the similarly minimal tetracycline molecule (0.03 µg/ml), substantially outperformed both eravacycline (0.25 µg/ml) and tigecycline (1 µg/ml).

**Figure 4:**
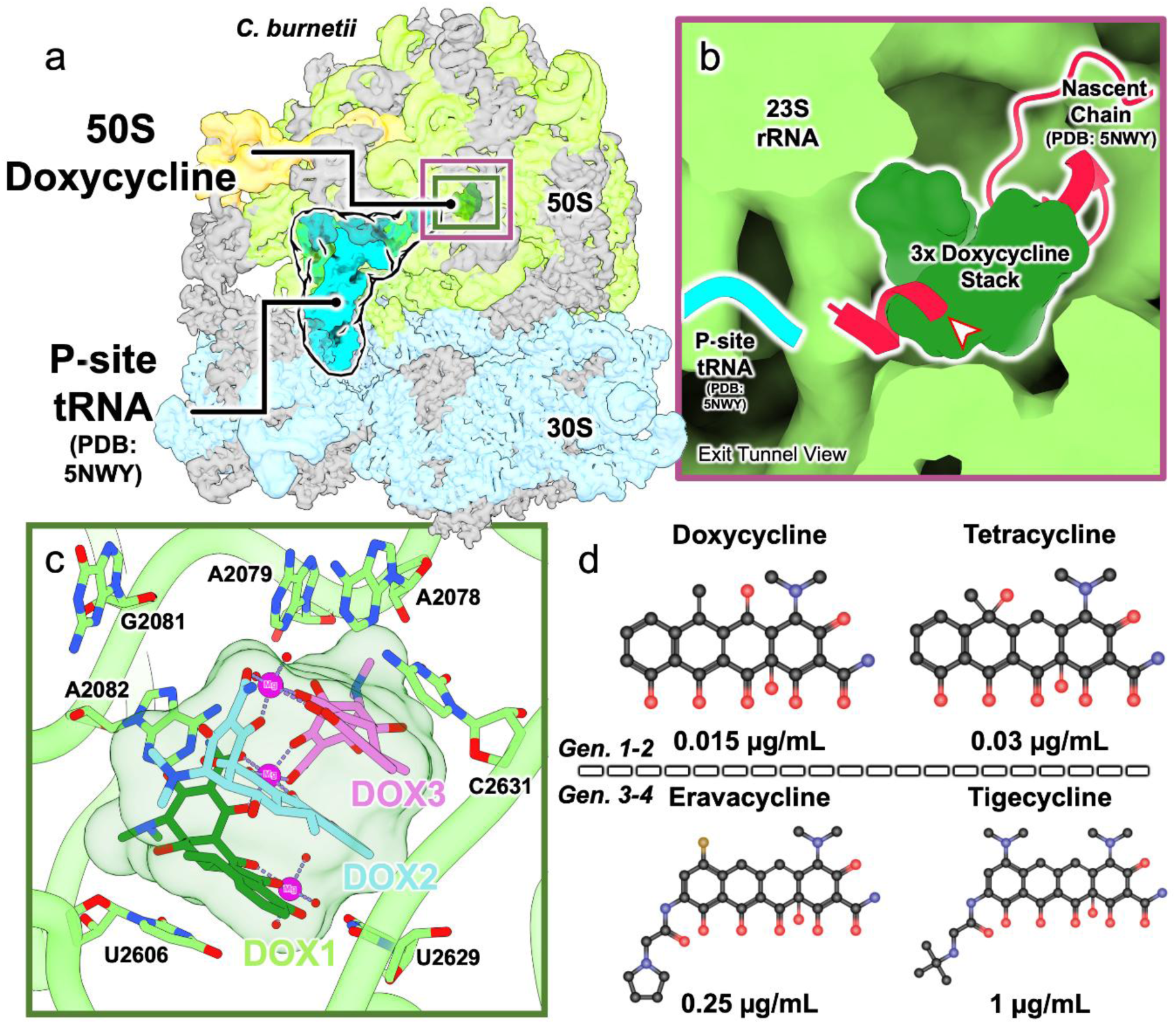
Three doxycycline molecules stack within the *C. burnetii* exit tunnel. (**a**) Overview of the doxycycline bound *C. burnetii* ribosome. All molecules shown as surface. Proteins are coloured grey, whilst r16S and r23S rRNA are coloured blue and green respectively. P-site tRNA (pdb_00005nwy, [92]) is shown in cyan blue and a novel stack of three doxycycline molecules in dark green (sphere atom representation). (**b**) Superposition of the two structures shows that doxycycline completely blocks the nascent chain channel. Cut-through image of the ribosome shown, with a nascent chain and P-site tRNA (both pdb_00005nwy, [92]) as cartoon representation in crimson and cyan respectively. (**c**) Three molecules of doxycycline form a stack between bases U2606 and C2631. rRNA shown as cartoon with interacting bases as sticks; doxycycline molecules shown as sticks coloured green, cyan, and magenta for molecules 1, 2, and 3 respectively. The doxycycline stack is coordinated around three magnesium ions (magenta spheres) that interact with doxycycline oxygen atoms and waters (not shown). (**d**) Later generation tetracyclines which display higher inhibitory values in other species perform worse than older, more compact chemicals. These may be unable to fit within the early exit tunnel in the same manner as doxycycline, suggesting the additional exit tunnel blocking mode of action explains the unusual potency for doxycycline against *C. burnetii*.

### A rearranged peptidyl transferase centre in the *E. coli* ribosome

Next, we attempted to identify whether doxycycline occupied the *E. coli* exit tunnel in a comparable way. 3D classification was carried out on the *E. coli* cryo-EM dataset, focusing on the *C. burnetii* doxycycline 50S binding site. The majority of particles showed the expected large subunit structure and weak density for a single doxycycline molecule (corresponding to DOX1; Fig. S1d). However, ∼12% of particles showed a markedly different structure. Refinement of these yielded a 2.16 Å structure showing a single bound doxycycline and major rearrangement of the tertiary structure of the PTC and early nascent chain exit tunnel between C2498-G2508 and G2056-C2064 (Fig. 5a-b, Fig. S11a). This rearrangement resembled a substantial ‘zippering’ of these regions of RNA (Movie 1). The early exit tunnel is typically lined with bases, however in this zippered structure these nucleotides are inverted so that the tunnel lumen is instead lined by phosphate backbone (Fig. S11). Doxycycline stacks on Ψ2504, with O3 and O21 anchoring the antibiotic via a magnesium to the phosphate of A2062, pinching both backbones together (Fig. 5a-b,g, Movie 1). The PTC is also now closed, preventing accommodation of tRNA in either the P or A-site (Fig. 5a-e) or nascent peptide. Notably, both C2506 (Fig. 5e) and U2063 (Fig. 5f) are splayed outwards, forming new interactions (Fig. S11c) and clashing with incoming P and A-site tRNAs. Other key rearrangements include A2062 ‘flipping in’, away from the PTC to base pair with deoxyuridine 2449 and, via a non-canonical base pair, with A2450. The A2450–C2063 base pair essential for translation is abolished [34]. Many additional new interactions are present, which we describe further in the Supplementary Discussion. Density for ribosomal protein bL4 is also notably poorer in the rearranged form than the canonical structure, due to a movement of the backbone around A2060, preventing the extended loop from anchoring against the phosphate backbone (Fig. S11b).

**Figure 5:**
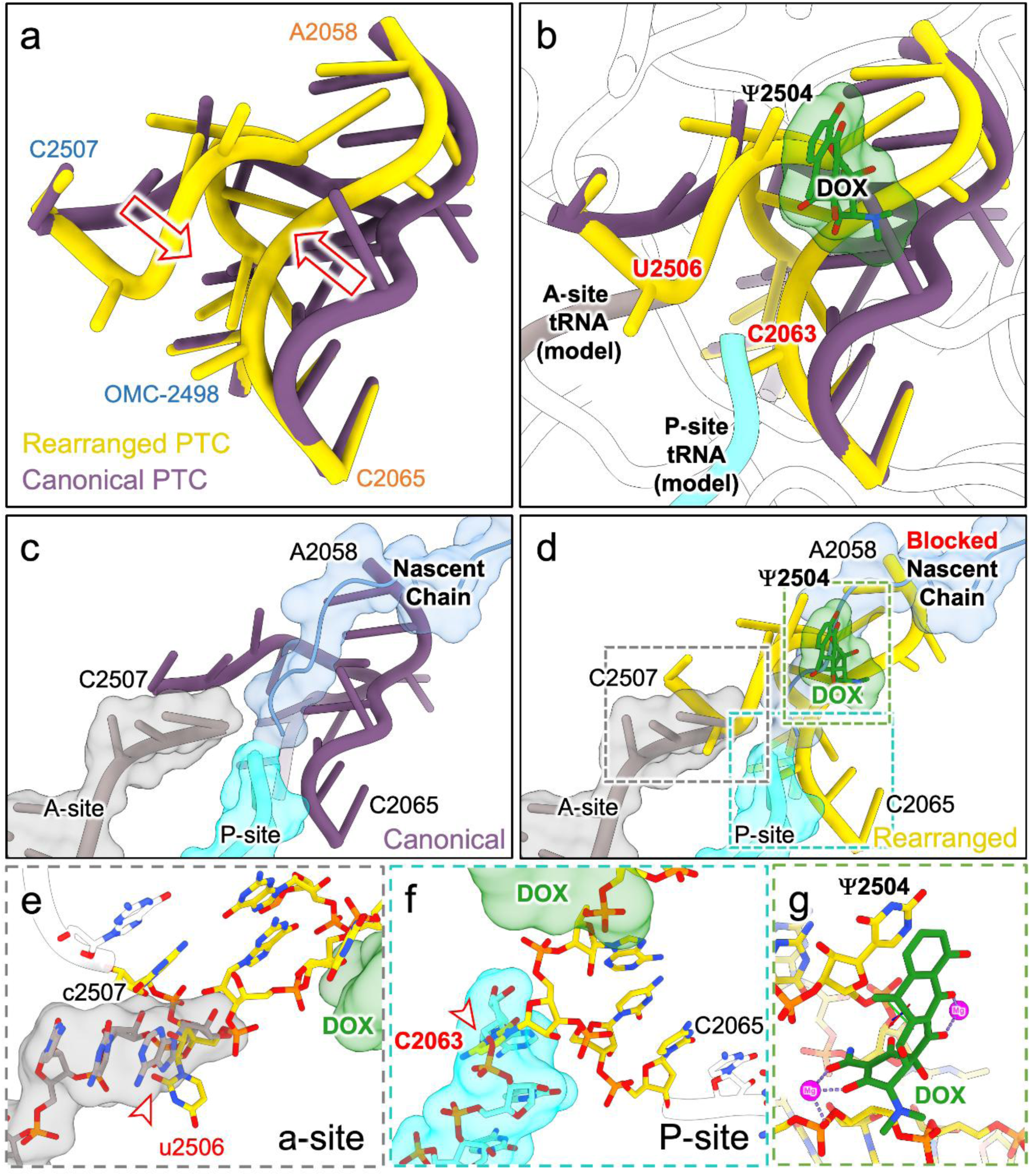
A rearranged peptidyl transferase centre. A subset of our *E. coli* data revealed a remarkable novel conformation of the prokaryotic ribosome bound to doxycycline. (**a**) Doxycycline drives a fundamental reorganisation of the r23S rRNA between bases 2058-2065 and 2498-2507 at the peptidyl transfer centre (PTC). rRNA is shown in cartoon representation, with the canonical structure in purple and the rearranged structure in yellow. The direction of movement is indicated by red arrows. (**b**) Doxycycline interacts with the rearranged PTC, stabilising this conformation that may be sampled at low occupancy in normal *E. coli* ribosomes. Doxycycline shown as sticks with transparent surface, carbon atoms, green; oxygen, red; nitrogen, blue. The position of the P-site tRNA from the active *E. coli* ribosome (pdb_00007k00, [73]) is shown as cyan cartoon. OMC: O2’-methylcytidine. (**c**) Bound doxycycline occupies the location typically held by the nascent chain (blue cartoon and transparent surface) in the canonical, translating structure (PDB ID: pdb_00007k00, [73]). (**d**) The rearranged PTC rRNA clashes with both A (mauve cartoon and transparent surface) and P-site tRNAs. This rearrangement will therefore prevent tRNA binding, peptide transfer, and the exit of a nascent chain. (**e**) The A-site tRNA is blocked by base U2506 which kinks outward from the exit tunnel in the rearranged PTC. rRNA and A-site tRNA shown as sticks; rRNA carbon, yellow, A-site tRNA carbon, mauve, phosphate, orange. (**f**) The P-site tRNA is blocked by base C2063 flipping outward from the exit tunnel. tRNA shown as sticks with transparent surface, carbon and surface cyan. (**g**) Detail of doxycycline interaction with rRNA. Doxycycline coordinates two magnesium ions with O11-O12 and O3-O21 respectively. Ring ‘D’ stacks upon pseudouridine Ψ2504.

## Discussion

HPF_cold_ represents the first structure of a hibernation factor with a cold shock domain, revealing a previously undetected member of the prokaryotic ribosome hibernation factor family [35]. Homologous cold shock domains are present across a wide range of bacterial phyla, particularly the proteobacteria, suggesting either an ancient origin with subsequent losses or horizontal gene transfer. Since it is structurally distinct from existing HPF family members (HPF_long_, HPF_short_, ribosome associated inhibitor A, RaiA; Fig. S2c) and the recently discovered Balon hibernation factor [35, 36] we propose to name this class HPF_cold_. HPF_cold_, like other members of the HPF family, occupies the A-site, blocking tRNA entry. HPF members differ in their effect on the ribosome, either stabilising a single ribosome (70S hibernation) or recruiting a partner ribosome to achieve ribosome dimers (100S hibernation) [35]. HPF_short_ functions in combination with ribosome modulation factor (RMF). Both RMF [37] and HPF_long_ [38] drive formation of 100S ribosome dimers that protect the mRNA entrance tunnel, particularly the 3’ end of the r16S rRNA, reducing susceptibility to RNAses, aiding survival, and increasing virulence [39]. *C. burnetii* lacks HPF_long_, RaiA, and RMF orthologues (its other hibernation factor (*CBU_0745*) having an HPF short structure, likely retained for shorter term hibernation) [37]. However, HPF_cold_ protects the 3’ end of the r16S rRNA by binding and stabilising the anti-Shine Dalgarno sequence through the CSD, blocking RNAse access to the hibernating ribosome. This likely confers a high enough degree of protection from RNAses to keep 70S ribosomes viable during the extended hibernation periods required of the *C. burnetii* SCV form [40].

We discovered a previously unknown peptide (CLaSP) closely associated with both the r23S and r5S rRNA. The role of CLaSP remains unclear, since the r5S rRNA of *C. burnetii* is truncated compared to other bacteria (Fig. S5a), suggesting a role in ribosome assembly and r23S and r5S stability [41, 42]. The position of CLaSP overlaps that of the mycobacterial specific ribosomal protein bL37 (Fig. S5b) [43]. Given its proximity to the PTC, bL37 has been hypothesised to function in communication within the ribosome, a function CLaSP may also provide [32]. CLaSP does not share significant sequence identity or structural characteristics with bL37. Indeed, superposing the structures indicates that a CLaSP-like peptide would not bind to the r5S rRNA of *Mycobacteria*. CLaSP is structurally dissimilar to the uL30 extension from *Borrelia burgdorferi*, which occupies the same cleft [57]. Finally, it is possible that CLaSP may mitigate the general reduction in ribosomal PTMs we observe in *C. burnetii*, compared to *E. coli* (Fig. S1) [44, 45], increasing the efficiency of this reduced ribosome. The hypothesis that there has been pressure to eliminate unnecessary modifications is supported by the remaining *C. burnetii* rRNA modification genes mapping precisely to those previously identified as providing the most significant growth rescue in an *E. coli* multiple gene knockout [46].

The *C. burnetii* ribosomal exit tunnel in the 50S subunit is filled by a remarkable stack of three doxycycline molecules (Fig. 4a-c), blocking nascent peptide exit (as observed for other antibiotic classes [47]). To date, no antibiotic has been observed to block the exit tunnel so extensively. This stack stabilises U2506 (Fig. S9c) and A2082 (Fig. S9e), which sense ribosome stalling [48] and translation arrest [49] (Fig. S8 and S9; [50]). The tetracycline-like antimicrobials sarecycline and tetracenomycin X also bind to the exit tunnel [33, 47], in a similar area to DOX1 (Fig. S9). However, they are rotated with respect to DOX1 and so could not accommodate the three-molecule stack or magnesium coordination. This doxycycline triple stack therefore represents a novel mode of antibiotic action that has greater similarity to the macrolide mode of action (Supplementary Discussion).

A major driver for the development of later generation tetracyclines was maintaining or improving binding to the small subunit. Eravacycline has a ∼14 times greater binding affinity for the 30S subunit than tetracycline in *E. coli* [51]. The 30S binding site is present in the *C. burnetii* ribosome, however the smaller early generation tetracyclines show much greater potency against *C. burnetii.* This suggests that the binding in the large subunit, which can only accommodate the smaller tetracyclines, is the major determinant of the doxycycline MIC in this pathogen. The ability to bind both large and small subunits presents the opportunity for complementarity and synergy, increasing the barrier for resistance mutations. However, it should be noted that some strains reporting reduced sensitivity to doxycycline have been reported [52]. Our results suggest that the traditional focus on optimising tetracycline binding to the small subunit may not improve antimicrobial activity universally.

Unexpectedly, we observed doxycycline bound to a major rearrangement of the *E. coli* peptidyl transferase centre (Fig. 5). To the best of our knowledge, the ribosome has never previously been observed in this state, nor has an antibiotic been observed to trigger such a major structural transformation. The peptidyl transferase centre has been thought to be relatively static, with little backbone conformational variation between ribosomal states or between the domains of life [53–57]. The structure we have captured challenges this, revealing that the prokaryotic ribosome is capable of a significant rearrangement within this extremely conserved, ancient region. Of particular note is the abolition of the essential A2450-C2063 interaction. Although we observed this conformation in only a subset of purified ribosome particles, it is possible that it is sampled more frequently in the cell than in purified ribosomes. The altered conformation will block the tip of the tRNA acceptor stem from the P and A-sites and would be unable to accommodate the nascent chain (Fig. 5c-f, Movie 1). The other major change is disordering of residues 60-65 of the bL4 loop (Fig. S10b). Deletion of this loop has previously been shown to slow translation and disrupt ribosome formation [58]. Alterations of the bL4 loop negatively impact translation fidelity and confer erythromycin resistance [59, 60]. Evidently, this ribosome conformation will be severely impaired and is likely inactive. This suggests an explanation for the bactericidal properties of doxycycline observed against *E. coli* strain ED1a, which shows particular bactericidal sensitivity to doxycycline [61]. In another recent study, ED1a did not show a lowered MIC value against tetracycline compared to other *E. coli* strains, but bactericidal activity remained, appearing to arise from an effect against the ribosome itself [62]. Thermal proteomic profiling results comparing *E. coli* ED1a (bactericidal) and *E. coli* K12 (bacteriostatic) showed the main divergence arising from the large subunit of ED1a ribosomes. While large subunit binding was not identified in that study, the classification required of our data suggests it could occur in a subset of ribosomes, making it difficult to identify. This inactive conformation of the ribosome presents an attractive avenue for future antibiotic development. Modification of doxycycline to exploit the properties of this site could lead to a potent inhibitor, potentially with bactericidal properties.

In conclusion, we have identified two previously unobserved binding modes of doxycycline to the ribosome large subunit in distinct bacterial species. One explains the proclivity of doxycycline for the *C. burnetii* ribosome while the other reveals a striking inactive ribosome conformation in *E. coli*. The modes of inhibition are mutually exclusive – one fills the nascent chain exit tunnel and the other remodels it. In addition, we reveal a new *C. burnetii* gene encoding a ribosomal peptide and describe a new member of the prokaryotic hibernation factor family, HPF_cold._ In a climate of increasing antimicrobial resistance, these offer new routes for antibiotic development.

## Methods

### *E. coli* ribosome isolation

400 ml BL21 DE3 (Merck #69450-3) *Escherichia coli* cells were grown to OD_600_ 0.7 and isolated through centrifugation at 4,000 x*g* for 15 minutes at 20°C. Pelleted cells were resuspended in 7 ml filtered and autoclaved BS100 buffer containing: 25 mM HEPES pH 7.5, 100 mM KOAc, 15 mM Mg(OAc)_2_, 1 mM DTT (added prior to each use) and frozen at −20°C.

Thawed cells were lysed in a Precellys 24 homogeniser, utilising VK01 2 ml tubes with 0.1 mm zircon beads. Samples were lysed with 8x 20 s beating at 6,500 rpm and 5 minutes of cooling on ice between each run. Supernatant was collected and beads were washed with buffer to elute additional lysate. Cell debris were subsequently clarified by centrifugation at 9,000 x*g* for 15 minutes at 4°C.

Ribosome purification followed previously described methods [63]. Supernatant was layered over a 30% (w/v) sucrose in BS100 cushion and centrifuged at 206,000 x*g* for 3 hours at 4°C. Ribosome pellet was resuspended in 500 µL BS100 buffer through shaking at 450 rpm at 4°C for 90 minutes using a Stuart Scientific SA8 vortex mixer (Merck #Z648531). Suspension was layered over a 10-40% (w/v) BS100 sucrose gradient and centrifuged at 125,000 x*g* for 80 min at 4°C. Cushion was manually aspirated into 500 µL fractions and 70S fractions were pooled and concentrated in a 100 kDa Vivaspin concentrator (Sartorius VS0142). Purified samples were snap frozen in liquid nitrogen and stored at −80 °C.

### *E. coli* ribosome grid preparation

Purified ribosome samples were diluted to approximately 0.3 mg/ml (112.5 nM) and incubated with a final concentration of 50 µM doxycycline hyclate (Merck #D9891) for one hour on ice. 3 µL ribosome antibiotic complex was added to a Quantifoil R2/2 2 nm continuous carbon coated grid (Agar Scientific, AGS173-2-2CL), blotted for 4 s at 4°C, 100% humidity before plunge freezing with a Vitrobot Mark IV (ThermoFisher Scientific).

Electron micrographs were collected at the electron biological imaging centre (eBIC) using Krios I (FEI Titan Krios, ThermoFisher) with a K3 detector (Gatan). 34,720 micrographs were collected at 130 k magnification with a pixel size of 0.615 Å and a total exposure of 45 e^-^/Å^2^.

### *E. coli* doxycycline bound structure data processing

Raw data were imported into cryoSPARC (v4) for patch motion correction and CTF estimation. 2,556 low quality micrographs (CTF fit greater than 5 Å) were removed. Data were initially processed in two batches in parallel. 774k particles were picked from Batch 1 (17,587 micrographs) and subject to two rounds of 2D classification (4x binning, 2.6 Å/pix). The resulting 417k particles were subject to ab-initio classification. 392k particles were re-extracted with 2x binning (1.3 Å/pix) yielding a reconstruction at the Nyquist limit (2.6 Å). Batch 2 (14,523 micrographs) blob picking yielded 1.46m particles, reducing to 967,764 following one round of 2D classification. All blob picked particles were then pooled and combined with crYOLO general model particle picks, subject to duplicate removal and subject to further 2D classification. Ab-initio and heterogenous refinement were used to assess particle quality and select for high quality 70S ribosomes. This yielded 1.7 million particles which were re-extracted at 0.8 Å/pix. Homogenous refinement yielded a 70S ribosome at 2.11 Å, improving to 1.95 Å using non-uniform refinement. Reference-based motion correction followed by non-uniform refinement with global and local CTF-refinement resulted in a 1.77 Å 70S reconstruction. Local refinement of the 30S subunit with the same particle set yielded a 1.94 Å reconstruction.

Classification was carried out on the hibernation factor binding site to separate doxycycline and YfiA bound ribosomes. A mask around YfiA was created and 3D classification without alignment was used to classify based on occupancy. 1.07m particles were in the YfiA bound class and 677k in the doxycycline/tRNA bound class. A local refinement of the YfiA class gave a map refined to 1.94 Å. A repeated classification (same parameters) on the 677k doxycycline/tRNA class selected 306k particles bound to doxycycline without tRNA and 372k with both present. Local refinements resolved to 2.13 Å without tRNA and 2.08 Å with tRNA however antibiotic density was more easily interpretable in the doxycycline with tRNA class, so this refinement was utilised for model building.

Following identification of the 50S doxycycline binding in *C. burnetii* focused classification was performed on the peptidyl transferase centre in *E. coli*. This utilised a mask around the three antibiotic molecules and 10 classes with a high-resolution filter (3 Å). All Fourier shell correlation (FSC) curves shown in Fig. S13.

### *C. burnetii* cell lysis

mCherry-NMII *C. burnetii* cells [64] (inoculated with 100 µL 1×10^7^ stock) were grown in 4×50 ml 1x ACCM-2 (Sunrise Bioscience #4700-300) in Nunc 50 ml flasks (Thermo Scientific, #156367) under 2.5 % O_2_, 5% CO_2_ (referred to subsequently as low oxygen) at 37°C [65]. After 7 days of growth cells were harvested into 50 ml falcon tubes and pelleted by centrifugation at 8,000 x*g* for 20 minutes at room temperature. Pellets were resuspended in BS100 and pooled to a final volume of ∼8 ml before storage at −80°C. Cell lysis was carried out using bead beating (Precellys 24, 8×1 ml added to VK01 beads, Bertin Corp.) to eliminate the risk of aerosolisation that can arise with other lysis methods. Lysis protocol carried out as described above with *E. coli* except each 1 ml lysate sample was kept separate for a subsequent sterility check (below). Lysate was subject to centrifugation at 18,000 x*g* for one hour to pellet any remaining cells and filtered through a 0.22 µM filter. Sample not used for sterility test was split into 100 microlitre aliquots in 0.5 ml Eppendorf tubes, snap frozen in liquid nitrogen and stored at −80°C.

### *C. burnetii* sterility check

100 microlitres of each lysate fraction was inoculated into 5 ml ACCM-2 and incubated statically for 7 days at 37°C under low oxygen. A positive (20 microlitres at 2×10^10^ cfu/ml, −80 °C stock) and negative control (100 microlitres ACCM-2 alone) were used. After 7 days, ACCM-2 0.25% (w/v) agar plates were inoculated with 250 microlitres culture, incubated for 10 days at 37°C under low oxygen and assessed for growth.

### *C. burnetii* ribosome isolation

Post-sterility check pooled lysate was layered onto a 30% (w/v) sucrose BS100 cushion and centrifuged at 206,000 x*g* for 3 hours at 4°C. The ribosome pellet was resuspended in 500 µL BS100 buffer supplemented with 300 nM doxycycline, shaking at 450 rpm at 4°C for 90 minutes as for *E. coli*. Ribosomes were concentrated to 0.05 mg/ml using a 100 kDa Vivaspin concentrator (Sartorius VS0142) and snap frozen in liquid nitrogen for −80 °C storage.

### *C. burnetii* ribosome grid preparation and data collection

Ribosomes were incubated with doxycycline hyclate (Merck #D9891) at a final concentration of 50 µM for one hour on ice. Grids were frozen as for *E. coli*.

Electron micrographs were collected at the electron biological imaging centre (eBIC) using Krios IV (FEI Titan Krios) with a K3 detector (Gatan). 22,295 micrographs were collected at 81 k magnification with a pixel size of 1.06 Å and a total exposure of 45 e^-^/Å^2^.

### *C. burnetii* doxycycline bound structure data processing

Raw data were imported into cryoSPARC (v4) for patch motion correction and CTF estimation [66–68]. A crYOLO convolutional neural network model was trained with blob-picked particles cleaned using 2D classification [69]. 5.54m crYOLO picked particles were extracted with 4x binning and subject to four rounds of 2D classification, resulting in 305k particles. One round of ab-initio 3D classification and a final 2D classification yielded 222k final particles for refinement. Particles were un-binned and refined to 2.71 Å with homogenous refinement, improving to 2.40 Å with non-uniform refinement [70]. Particles were separated by exposure group and subject to global CTF refinement followed by reference-based motion correction. Non-uniform refinement gave a final 50S reconstruction to 2.15 Å.

Focused classification on the 50S doxycycline binding site was carried out as above for *E. coli*. This resulted in 91k 50S antibiotic bound and 129k empty ribosomes. These were refined to 2.22 Å and 2.19 Å respectively using non-uniform refinement. Despite further classification on the 50S, an *E. coli-*like rearranged ribosome class could not be identified.

Focused 3D classification without alignment isolated 44k 70S ribosome particles, giving a 2.48 Å 70S non-uniform reconstruction and a 2.83 Å local refinement of the 30S subunit. Focused classification without alignment was carried out with a mask around HPF_cold_, as for *E. coli*. This identified 24k HPF_cold_ bound ribosomes and 20k 30S doxycycline and tRNA bound ribosomes. The HPF_cold_ bound ribosome refined to 2.87 Å with local refinement. As for *E. coli*, a further round of classification of the doxycycline bound particles removed non-tRNA bound ribosomes which displayed poorer quality density. This resulted in a 3.02 Å local refinement of doxycycline bound to the 30S from 11k particles. Fourier shell correlation curves shown in Fig. S13.

### Atomic model building

*E. coli* model building was carried out in both Coot and Isolde [71, 72] using the published 2 Å *E. coli* structure (PDB 7k00) as a starting model [73]. Waters and magnesium ions were manually checked and built where necessary. Final refinements were carried out with Refmac v5.8 [74] and Phenix v1.21 [75].

*C. burnetii* AlphaFold v2.2 [76] ribosomal protein predictions were rigid body fitted into electron density using ChimeraX v1.7 [77] and rebuilt using Coot and Isolde [71, 72]. ModelAngelo [78] was used to provide an initial template for rRNA which was rebuilt and sequence corrected as necessary. r5S rRNA and L1 stalk AlphaFold 3 [79] predictions were incorporated into the structure and rebuilt. Where it was not possible to confidently identify an rRNA post-transcriptional modification from the cryo-EM map, we included it only if the genome encoded a corresponding post-transcriptional modification protein. Final models were refined using Refmac and Phenix [74, 75].

### *C. burnetii* minimum inhibitory concentration

Doxycycline and tetracycline stocks were both reconstituted in dH_2_O at 10 mg/ml. Eravacycline (CAS 1334714-66-7) and tigecycline (CAS 220620-09-7) were purchased from Cambridge Bioscience. Eravacycline was reconstituted in distilled H_2_O to 1 mg/ml and tigecycline reconstituted in dimethyl sulfoxide (DMSO) to 10 mg/ml (a DMSO serial dilution control was included). All antibiotics were stored at −20 °C. Antibiotic serial dilutions were carried out in ACCM-2 and inoculated with final 1×10^6^ *C. burnetii* mCherry-NMII cells. After 10 days or with clear growth in positive controls, MIC values were determined as described previously [80]. All MIC assays were conducted with a minimum of three repeats.

### HPF_cold_ phylogenetic analysis

Protein sequences containing both PF02482 and PF00313 Pfam domains were retrieved from UniProtKB1 [81]. Domains were identified with hmmsearch, part of HMMER v3.3.2 software suite2 [82], against the corresponding Pfam HMMs, using Pfam gathering thresholds3. Only sequences with significant matches to both target domains were retained. To remove exact duplicates, sequences were clustered with CD-HIT v4.8.14 at 100% identity [83]. Multiple sequence alignment was generated with MAFFT v7.4905 using the G-INS-i algorithm (global-homology strategy) [84]. The alignment was trimmed with trimAl v1.4.rev156 using the gappyout procedure to remove poorly aligned and gap-rich positions while preserving informative sites [85]. Maximum-likelihood trees were reconstructed with IQ-TREE v2.2.07 [86] with the best-fit substitution model selected by ModelFinder8 under the Bayesian Information Criterion (BIC) [87]. Branch support was assessed using ultrafast bootstrap9 (10,000 replicates). Trees were visualised and annotated in iTOL v6 [88].

## Acknowledgements

We acknowledge Diamond for access and support of the cryo-EM facilities at the UK national electron Bio-Imaging Centre (eBIC), proposal BI25452-50 (*E. coli* data collection) and BI32707-9 (*C. burnetii* data collection). We also thank the staff of eBIC for collection of the cryo-EM data, in particular Karen Davies and her in-person assistance collecting on Krios I. We thank Becky Conners (University of Exeter) for guidance with ribosome purification. We also thank Ufuk Borucu for guidance in preparing samples for data collection and acknowledge the GW4 Facility for High-Resolution Electron Cryo-Microscopy, funded by the Wellcome Trust (202904/Z/16/Z and 206181/Z/17/Z) and BBSRC (BB/R000484/1). We thank Prof. Robert Heinzen (Rocky Mountain Laboratories, MT) for providing the mCherry NMII *C. burnetii* strain. WSS was supported by the BBSRC (studentship BB/M009122/1) and Dstl (contract DSTLX1000167246). BD and MM were supported by an ERC Starting Grant under the European Union’s Horizon 2020 research and innovation program (grant agreement No 803894), awarded to BD. BD also received funding from the UK Research and Innovation (UKRI) under the UK government’s Horizon Europe funding guarantee (EP/Y037049/1), as well as the the Leverhulme Trust (grant reference RPG-2024-417). VG received funding from the BBSRC (grant number BB/R008639/1) and Leverhulme Trust (grant reference RPG-2023-069). For the purpose of open access, the author has applied a Creative Commons Attribution (CC BY) license to any Author Accepted Manuscript version arising from this submission.

## Data availability

The structures associated with this manuscript have been submitted to the Protein Data Bank with the codes: 9SLG, 9ST6, 9SX2, 9T0Z, 9T1N, 9TY2, and 9T61. The cryo-EM data have been submitted to the Electron Microscopy Data Bank (EMDB) with the codes EMD-55001, EMD-55212, EMD-55330, EMD-55416, EMD-55439, EMD-55479, and EMD-55601.

## References

1. Celina, S.S. and J. Cerny, Coxiella burnetii in ticks, livestock, pets and wildlife: A mini-review. Front Vet Sci, 2022. 9: p. 1068129.

2. van Schaik, E.J., et al., Molecular pathogenesis of the obligate intracellular bacterium Coxiella burnetii. Nat Rev Microbiol, 2013. 11(8): p. 561–73.

3. Neare, K., et al., Coxiella burnetii Antibody Prevalence and Risk Factors of Infection in the Human Population of Estonia. Microorganisms, 2019. 7(12): p. 629.

4. Brooke, R.J., et al., Exposure to low doses of Coxiella burnetii caused high illness attack rates: Insights from combining human challenge and outbreak data. Epidemics, 2015. 11: p. 1–6.

5. Brooke, R.J., et al., Human dose response relation for airborne exposure to Coxiella burnetii. BMC Infect Dis, 2013. 13: p. 488.

6. Wiebe, M.E., P.R. Burton, and D.M. Shankel, Isolation and characterization of two cell types of Coxiella burneti phase I. J Bacteriol, 1972. 110(1): p. 368–77.

7. Angelakis, E. and D. Raoult, Q Fever. Vet Microbiol, 2010. 140(3-4): p. 297–309.

8. Cross, A.R., et al., Zoonoses under our noses. Microbes Infect, 2019. 21(1): p. 10–19.

9. Schneeberger, P.M., et al., Q fever in the Netherlands - 2007-2010: what we learned from the largest outbreak ever. Med Mal Infect, 2014. 44(8): p. 339–53.

10. Morroy, G., et al., Of goats and humans; the societal costs of the Dutch Q fever saga. International Journal of Infectious Diseases, 2012. 16: p. e266.

11. Honarmand, H., Q Fever: an old but still a poorly understood disease. Interdiscip Perspect Infect Dis, 2012. 2012: p. 131932.

12. Buijs, S.B., et al., Still New Chronic Q Fever Cases Diagnosed 8 Years After a Large Q Fever Outbreak. Clin Infect Dis, 2021. 73(8): p. 1476–1483.

13. UKHSA, UKHSA highlights pathogens of greatest risk to public health. 2025. UKHSA highlights pathogens of greatest risk to public health - GOV.UK, cited 23rd October 2025 (2025).

14. Clemente, T.M., R.K. Angara, and S.D. Gilk, Establishing the intracellular niche of obligate intracellular vacuolar pathogens. Front Cell Infect Microbiol, 2023. 13: p. 1206037.

15. Kersh, G.J., Antimicrobial therapies for Q fever. Expert Rev Anti Infect Ther, 2013. 11(11): p. 1207–14.

16. Kazar, J., Coxiella burnetii infection. Ann N Y Acad Sci, 2005. 1063: p. 105–14.

17. Maurin, M., et al., Phagolysosomal alkalinization and the bactericidal effect of antibiotics: the Coxiella burnetii paradigm. J Infect Dis, 1992. 166(5): p. 1097–102.

18. Clay, K.A., et al., Evaluation of the Efficacy of Doxycycline, Ciprofloxacin, Levofloxacin, and Co-trimoxazole Using In Vitro and In Vivo Models of Q Fever. Antimicrob Agents Chemother, 2021. 65(11): p. e0067321.

19. Cunha, B.A., P. Domenico, and C.B. Cunha, Pharmacodynamics of doxycycline. Clin Microbiol Infect, 2000. 6(5): p. 270–3.

20. Agafonov, D.E., V.A. Kolb, and A.S. Spirin, Ribosome-associated protein that inhibits translation at the aminoacyl-tRNA binding stage. EMBO Rep, 2001. 2(5): p. 399–402.

21. Prossliner, T., et al., Hibernation factors directly block ribonucleases from entering the ribosome in response to starvation. Nucleic Acids Res, 2021. 49(4): p. 2226–2239.

22. Ekemezie, C.L. and S.V. Melnikov, Hibernating ribosomes as drug targets? Front Microbiol, 2024. 15: p. 1436579.

23. Park, D., et al., Developmental Transitions Coordinate Assembly of the Coxiella burnetii Dot/Icm Type IV Secretion System. Infect Immun, 2022. 90(10): p. e0041022.

24. Brodersen, D.E., et al., The structural basis for the action of the antibiotics tetracycline, pactamycin, and hygromycin B on the 30S ribosomal subunit. Cell, 2000. 103(7): p. 1143–54.

25. Pioletti, M., et al., Crystal structures of complexes of the small ribosomal subunit with tetracycline, edeine and IF3. EMBO J, 2001. 20(8): p. 1829–39.

26. Jenner, L., et al., Structural basis for potent inhibitory activity of the antibiotic tigecycline during protein synthesis. Proc Natl Acad Sci U S A, 2013. 110(10): p. 3812–6.

27. Afseth, G., Y.Y. Mo, and L.P. Mallavia, Characterization of the 23S and 5S rRNA genes of Coxiella burnetii and identification of an intervening sequence within the 23S rRNA gene. J Bacteriol, 1995. 177(10): p. 2946–9.

28. Warrier, I., et al., The Intervening Sequence of Coxiella burnetii: Characterization and Evolution. Front Cell Infect Microbiol, 2016. 6: p. 83.

29. Berman, H.M., et al., The Protein Data Bank. Nucleic Acids Res, 2000. 28(1): p. 235–42.

30. Altschul, S.F., et al., Basic local alignment search tool. J Mol Biol, 1990. 215(3): p. 403–10.

31. Mueller, U., et al., Thermal stability and atomic-resolution crystal structure of the Bacillus caldolyticus cold shock protein. J Mol Biol, 2000. 297(4): p. 975–88.

32. van Kempen, M., et al., Fast and accurate protein structure search with Foldseek. Nat Biotechnol, 2024. 42(2): p. 243–246.

33. Lomakin, I.B., et al., Sarecycline inhibits protein translation in Cutibacterium acnes 70S ribosome using a two-site mechanism. Nucleic Acids Res, 2023. 51(6): p. 2915–2930.

34. Chirkova, A., et al., The role of the universally conserved A2450-C2063 base pair in the ribosomal peptidyl transferase center. Nucleic Acids Res, 2010. 38(14): p. 4844–55.

35. Prossliner, T., et al., Ribosome Hibernation. Annu Rev Genet, 2018. 52: p. 321–348.

36. Helena-Bueno, K., et al., A new family of bacterial ribosome hibernation factors. Nature, 2024. 626(8001): p. 1125–1132.

37. Polikanov, Y.S., G.M. Blaha, and T.A. Steitz, How hibernation factors RMF, HPF, and YfiA turn off protein synthesis. Science, 2012. 336(6083): p. 915–8.

38. Franken, L.E., et al., A general mechanism of ribosome dimerization revealed by single-particle cryo-electron microscopy. Nat Commun, 2017. 8(1): p. 722.

39. Gohara, D.W. and M.F. Yap, Survival of the drowsiest: the hibernating 100S ribosome in bacterial stress management. Curr Genet, 2018. 64(4): p. 753–760.

40. Kersh, G.J., et al., Presence and persistence of Coxiella burnetii in the environments of goat farms associated with a Q fever outbreak. Appl Environ Microbiol, 2013. 79(5): p. 1697–703.

41. Bubunenko, M., et al., Nus transcription elongation factors and RNase III modulate small ribosome subunit biogenesis in Escherichia coli. Mol Microbiol, 2013. 87(2): p. 382–93.

42. Singh, N., et al., SuhB Associates with Nus Factors To Facilitate 30S Ribosome Biogenesis in Escherichia coli. mBio, 2016. 7(2): p. e00114.

43. Hentschel, J., et al., The Complete Structure of the Mycobacterium smegmatis 70S Ribosome. Cell Rep, 2017. 20(1): p. 149–160.

44. Ofengand, J. and M. Del Campo, Modified Nucleosides of Escherichia coli Ribosomal RNA. EcoSal Plus, 2004. 1(1): p. 4.6.1.

45. Fleming, A.M., et al., Direct Nanopore Sequencing for the 17 RNA Modification Types in 36 Locations in the E. coli Ribosome Enables Monitoring of Stress-Dependent Changes. ACS Chem Biol, 2023. 18(10): p. 2211–2223.

46. Ero, R., et al., Ribosomal RNA modification enzymes stimulate large ribosome subunit assembly in E. coli. Nucleic Acids Res, 2024. 52(11): p. 6614–6628.

47. Osterman, I.A., et al., Tetracenomycin X inhibits translation by binding within the ribosomal exit tunnel. Nat Chem Biol, 2020. 16(10): p. 1071–1077.

48. Koch, M., et al., Critical 23S rRNA interactions for macrolide-dependent ribosome stalling on the ErmCL nascent peptide chain. Nucleic Acids Res, 2017. 45(11): p. 6717–6728.

49. Vazquez-Laslop, N., et al., The key function of a conserved and modified rRNA residue in the ribosomal response to the nascent peptide. EMBO J, 2010. 29(18): p. 3108–17.

50. Santos, O.M., Silva, D.M., Martins, F.T., Legendre, A.O., Azarias, L.C., Rosa, I.M., Neves, P.P., de Araujo, M.B. and Doriguetto, A.C., Protonation pattern, tautomerism, conformerism, and physicochemical analysis in new crystal forms of the antibiotic doxycycline. Crystal growth & design, 2014. 14(8): p. 3711–3726.

51. Grossman, T.H., et al., Target- and resistance-based mechanistic studies with TP-434, a novel fluorocycline antibiotic. Antimicrob Agents Chemother, 2012. 56(5): p. 2559–64.

52. Rolain, J.M., F. Lambert, and D. Raoult, Activity of telithromycin against thirteen new isolates of C. burnetii including three resistant to doxycycline. Ann N Y Acad Sci, 2005. 1063: p. 252–6.

53. Ban, N., et al., The complete atomic structure of the large ribosomal subunit at 2.4 A resolution. Science, 2000. 289(5481): p. 905–20.

54. Schuwirth, B.S., et al., Structures of the bacterial ribosome at 3.5 A resolution. Science, 2005. 310(5749): p. 827–34.

55. Harms, J., et al., High resolution structure of the large ribosomal subunit from a mesophilic eubacterium. Cell, 2001. 107(5): p. 679–88.

56. Tirumalai, M.R., et al., The Peptidyl Transferase Center: a Window to the Past. Microbiol Mol Biol Rev, 2021. 85(4): p. e0010421.

57. Noller, H.F., et al., Structure of the ribosome at 5.5 A resolution and its interactions with functional ligands. Cold Spring Harb Symp Quant Biol, 2001. 66: p. 57–66.

58. Lawrence, M.G., et al., The extended loops of ribosomal proteins uL4 and uL22 of Escherichia coli contribute to ribosome assembly and protein translation. Nucleic Acids Res, 2016. 44(12): p. 5798–810.

59. O’Connor, M., S.T. Gregory, and A.E. Dahlberg, Multiple defects in translation associated with altered ribosomal protein L4. Nucleic Acids Res, 2004. 32(19): p. 5750–6.

60. Gregory, S.T. and A.E. Dahlberg, Erythromycin resistance mutations in ribosomal proteins L22 and L4 perturb the higher order structure of 23 S ribosomal RNA. J Mol Biol, 1999. 289(4): p. 827–34.

61. Maier, L., et al., Unravelling the collateral damage of antibiotics on gut bacteria. Nature, 2021. 599(7883): p. 120–124.

62. Khusainov, I., et al., Bactericidal effect of tetracycline in E. coli strain ED1a may be associated with ribosome dysfunction. Nat Commun, 2024. 15(1): p. 4783.

63. McLaren, M., et al., CryoEM reveals that ribosomes in microsporidian spores are locked in a dimeric hibernating state. Nat Microbiol, 2023. 8(10): p. 1834–1845.

64. Beare, P.A., et al., Characterization of a Coxiella burnetii ftsZ mutant generated by Himar1 transposon mutagenesis. J Bacteriol, 2009. 191(5): p. 1369–81.

65. Omsland, A., et al., Isolation from animal tissue and genetic transformation of Coxiella burnetii are facilitated by an improved axenic growth medium. Appl Environ Microbiol, 2011. 77(11): p. 3720–5.

66. Punjani, A., et al., cryoSPARC: algorithms for rapid unsupervised cryo-EM structure determination. Nat Methods, 2017. 14(3): p. 290–296.

67. Zivanov, J., T. Nakane, and S.H.W. Scheres, Estimation of high-order aberrations and anisotropic magnification from cryo-EM data sets in RELION-3.1. IUCrJ, 2020. 7(Pt 2): p. 253–267.

68. Rubinstein, J.L. and M.A. Brubaker, Alignment of cryo-EM movies of individual particles by optimization of image translations. J Struct Biol, 2015. 192(2): p. 188–95.

69. Wagner, T., et al., SPHIRE-crYOLO is a fast and accurate fully automated particle picker for cryo-EM. Commun Biol, 2019. 2: p. 218.

70. Punjani, A., H. Zhang, and D.J. Fleet, Non-uniform refinement: adaptive regularization improves single-particle cryo-EM reconstruction. Nat Methods, 2020. 17(12): p. 1214–1221.

71. Emsley, P. and K. Cowtan, Coot: model-building tools for molecular graphics. Acta Crystallogr D Biol Crystallogr, 2004. 60(Pt 12 Pt 1): p. 2126–32.

72. Croll, T.I., ISOLDE: a physically realistic environment for model building into low-resolution electron-density maps. Acta Crystallogr D Struct Biol, 2018. 74(Pt 6): p. 519–530.

73. Watson, Z.L., et al., Structure of the bacterial ribosome at 2 A resolution. Elife, 2020. 9: p. e60482.

74. Murshudov, G.N., et al., REFMAC5 for the refinement of macromolecular crystal structures. Acta Crystallogr D Biol Crystallogr, 2011. 67(Pt 4): p. 355–67.

75. Liebschner, D., et al., Macromolecular structure determination using X-rays, neutrons and electrons: recent developments in Phenix. Acta Crystallogr D Struct Biol, 2019. 75(Pt 10): p. 861–877.

76. Jumper, J., et al., Highly accurate protein structure prediction with AlphaFold. Nature, 2021. 596(7873): p. 583–589.

77. Pettersen, E.F., et al., UCSF ChimeraX: Structure visualization for researchers, educators, and developers. Protein Sci, 2021. 30(1): p. 70–82.

78. Jamali, K., et al., Automated model building and protein identification in cryo-EM maps. Nature, 2024. 628(8007): p. 450–457.

79. Abramson, J., et al., Accurate structure prediction of biomolecular interactions with AlphaFold 3. Nature, 2024. 630(8016): p. 493–500.

80. Clay, K.A., et al., Use of axenic media to determine antibiotic efficacy against Coxiella burnetii. Int J Antimicrob Agents, 2018. 51(5): p. 806–808.

81. UniProt, C., UniProt: the Universal Protein Knowledgebase in 2025. Nucleic Acids Res, 2025. 53(D1): p. D609–D617.

82. Potter, S.C., et al., HMMER web server: 2018 update. Nucleic Acids Res, 2018. 46(W1): p. W200–W204.

83. Fu, L., et al., CD-HIT: accelerated for clustering the next-generation sequencing data. Bioinformatics, 2012. 28(23): p. 3150–2.

84. Katoh, K. and D.M. Standley, MAFFT multiple sequence alignment software version 7: improvements in performance and usability. Mol Biol Evol, 2013. 30(4): p. 772–80.

85. Capella-Gutierrez, S., J.M. Silla-Martinez, and T. Gabaldon, trimAl: a tool for automated alignment trimming in large-scale phylogenetic analyses. Bioinformatics, 2009. 25(15): p. 1972–3.

86. Minh, B.Q., et al., IQ-TREE 2: New Models and Efficient Methods for Phylogenetic Inference in the Genomic Era. Mol Biol Evol, 2020. 37(5): p. 1530–1534.

87. Kalyaanamoorthy, S., et al., ModelFinder: fast model selection for accurate phylogenetic estimates. Nat Methods, 2017. 14(6): p. 587–589.

88. Hoang, D.T., et al., UFBoot2: Improving the Ultrafast Bootstrap Approximation. Mol Biol Evol, 2018. 35(2): p. 518–522.

89. Rusu, A. and E.L. Buta, The Development of Third-Generation Tetracycline Antibiotics and New Perspectives. Pharmaceutics, 2021. 13(12): p. 783.

90. Ahn, M., et al., Modulating co-translational protein folding by rational design and ribosome engineering. Nat Commun, 2022. 13(1): p. 4243.

91. Letunic, I. and P. Bork, Interactive Tree of Life (iTOL) v6: recent updates to the phylogenetic tree display and annotation tool. Nucleic Acids Res, 2024. 52(W1): p. W78–W82.

92. Su, T., et al., The force-sensing peptide VemP employs extreme compaction and secondary structure formation to induce ribosomal stalling. Elife, 2017. 6: p. e25642.

