## Supplementary information for "Cryo-EM reveals multiple mechanisms of ribosome inhibition by doxycycline"

**Stuart et al.**

**Supplementary information**

**Contents:**

**Supplementary Tables**

**Table S1: List of ribosomal proteins modelled for the *C. burnetii ribosome*** **2**

**Table S2: Data collection and refinement parameters** **3**

**Supplementary Figures**

**Figure S1: Electron density demonstrates a novel doxycycline binding site in *C. burnetii*.** **4**

**Figure S2: rRNA modifications observed in the *C. burnetii* ribosome** **5**

**Figure S3: $HPF_{cold}$ binds the 30S subunit of the ribosome, protecting the anti-Shine Dalgarno sequence** **7**

**Figure S4: $HPF_{cold}$ is an additional hibernation promoting factor** **8**

**Figure S5: Alignment of $HPF_{cold}$ with paralogues of the two domains** **10**

**Figure S6: CLaSP binding in context** **12**

**Figure S7: *C. burnetii* data processing pipeline** **14**

**Figure S8: *E. coli* data processing pipeline** **16**

**Figure S9: Electron density for doxycycline bound in the exit tunnel, comparison to the empty exit tunnel** **18**

**Figure S10: Structural details of doxycycline within the *C. burnetii* exit tunnel** **20**

**Figure S11: Inactive *E. coli* ribosome identified through doxycycline driven serendipity** **22**

**Figure S12: Novel rRNA interactions are made in the doxycycline bound *E. coli* ribosome** **24**

**Figure S13: Cryo-EM map and model validation** **25**

**Supplementary discussion** **27**

**References** **32**

**Cryo-EM reveals multiple mechanisms of ribosome inhibition by doxycycline**

**Stuart et al.**

**Supplementary Table 1: List of ribosomal proteins modelled for the *C. burnetii ribosome.*** Ribosomal proteins are numbered by homology to the *E. coli* ribosome. Gene names are from the *C. burnetii* RSA493 (Nine Mile) genome.

| Large Subunit Proteins | Gene | Small Subunit Proteins | Gene |
|---|---|---|---|
| bL2 | *CBU_0241* | bS3 | *CBU_0244* |
| bL3 | *CBU_0238* | bS4 | *CBU_0262* |
| bL4 | *CBU_0239* | bS5 | *CBU_0255* |
| bL5 | *CBU_0250* | bS6 | *CBU_0864* |
| bL6 | *CBU_0253* | bS7 | *CBU_0234* |
| bL9 | *CBU_0867* | bS8 | *CBU_0252* |
| bL13 | *CBU_1749* | bS9 | *CBU_1748* |
| bL14 | *CBU_0248* | bS10 | *CBU_0237* |
| bL15 | *CBU_0257* | bS11 | *CBU_0261* |
| bL16 | *CBU_0245* | bS12 | *CBU_0233* |
| bL17 | *CBU_0264* | bS13 | *CBU_0260* |
| bL18 | *CBU_0254* | bS14 | *CBU_0251* |
| bL19 | *CBU_0442* | bS15 | *CBU_0851* |
| bL20 | *CBU_1323* | bS16 | *CBU_0445* |
| bL21 | *CBU_0385* | bS17 | *CBU_0247* |
| bL22 | *CBU_0243* | bS18 | *CBU_0865* |
| bL23 | *CBU_0240* | bS19 | *CBU_0242* |
| bL24 | *CBU_0249* | bS20 | *CBU_0389* |
| bL25 | *CBU_1840* | bS21 | *CBU_1593* |
| bL27 | *CBU_0386* | HPFcold | *CBU_0020* |
| bL28 | *CBU_0291* | | |
| bL29 | *CBU_0246* | | |
| bL30 | *CBU_0256* | | |
| bL32 | *CBU_0491* | | |
| bL34 | *CBU_1917* | | |
| bL35 | *CBU_1324* | | |
| bL36 | *CBU_2097* | | |
| CLaSP | - | | |

### Cryo-EM reveals multiple mechanisms of ribosome inhibition by doxycycline


#### Stuart et al.


#### Supplementary Table 2: Data collection and refinement parameters

| | *E. coli* Rearrange | *E. coli* 30S Doxycycline | *C. burnetii* HPFcold | *C. burnetii* 30S Doxycycline | *C. burnetii* 70S (50S doxycycline & 30S HPFcold) | *C. burnetii* 50S Doxycycline | *C. burnetii* 50S Empty |
|---|---|---|---|---|---|---|---|
| **Data Collection** | | | | | | | |
| Magnification | 130k | 130k | 81k | 81k | 81k | 81k | 81k |
| Voltage (kV) | 300 | 300 | 300 | 300 | 300 | 300 | 300 |
| Electron exposure (e-/A2) | 45 | 45 | 45 | 45 | 45 | 45 | 45 |
| Defocus range (μM) | -0.4 to -2.5 (0.3 steps) | -0.4 to -2.5 (0.3 steps) | -0.8 to -2.0 (0.3 steps) | -0.8 to -2.0 (0.3 steps) | -0.8 to -2.0 (0.3 steps) | -0.8 to -2.0 (0.3 steps) | -0.8 to -2.0 (0.3 steps) |
| Pixel size (Å) | 0.645 | 0.645 | 1.06 | 1.06 | 1.06 | 1.06 | 1.06 |
| Symmetry imposed | C1 | C1 | C1 | C1 | C1 | C1 | C1 |
| Final particle images (no.) | 215,222 | 372,420 | 24,143 | 10,803 | 44,196 | 91,218 | 129,328 |
| Map resolution (Å) | 2.16 | 2.08 | 2.87 | 3.06 | 2.48 | 2.22 | 2.19 |
| FSC threshold | 0.143 | 0.143 | 0.143 | 0.143 | 0.143 | 0.143 | 0.143 |
| **Refinement** | | | | | | | |
| Composition (#) | | | | | | | |
| Chains | 63 | 25 | 31 | 30 | 72 | 41 | 33 |
| Atoms | 258161 (Hydrogens: 107886) | 91121 (Hydrogens: 37008) | 88761 (Hydrogens: 36589) | 86019 (Hydrogens: 35219) | 238507 (Hydrogens: 96251) | 148990 (Hydrogens: 59287) | 87980 (Hydrogens: 0) |
| Residues | Protein: 5946 Nucleotide: 4557 | Protein: 2492 Nucleotide: 1534 | Protein: 2413 Nucleotide: 1535 | Protein: 2239 Nucleotide: 1535 | Protein: 5707 Nucleotide: 4525 | Protein: 3245 Nucleotide: 2990 | Protein: 3011 Nucleotide: 2990 |
| Water | 5341 | 1356 | 5 | 5 | 17 | 15 | 34 |
| Ligands | K: 77 | K: 4 | K: 20 | K: 34 | K: 28 | K: 8 | K: 108 |
| | ZN: 2 | ZN: 1 | ZN: 2 | ZN: 2 | ZN: 2 | ZN: 3 | ZN: 1 |
| | MG: 446 | MG: 137 | MG: 2 | MG: 2 | MG: 14 | MG: 12 | MG: 157 |
| | MS6: 1 | IAS: 1 | ZN: 1 | ZN: 1 | ZN: 4 | DXT: 3 | |
| | DXT: 1 | DXT: 1 | | DXT: 1 | DXT: 3 | | |
| | IAS: 1 | | | | | | |
| Bonds (RMSD) | | | | | | | |
| Length (Å) (# > 4 σ) | 0.008 (116) | 0.007 (24) | 0.014 (32) | 0.006 (2) | 0.009 (32) | 0.005 (0) | 0.006 (16) |
| Angles (°) (# > 4 σ) | 1.178 (470) | 1.113 (28) | 1.875 (559) | 1.223 (179) | 1.266 (562) | 0.709 (3) | 1.251 (310) |
| MolProbity score | 1.66 | 1.21 | 1.53 | 1.03 | 1.45 | 1.36 | 1.03 |
| Clash score | 11.47 | 2.35 | 4.77 | 1.78 | 4.61 | 4.28 | 1.32 |
| Ramachandran plot (%) | | | | | | | |
| Outliers | 0 | 0 | 0.13 | 0 | 0.05 | 0 | 0 |
| Allowed | 2.42 | 3.19 | 2.83 | 2 | 2.5 | 2.29 | 2.47 |
| Favored | 97.58 | 96.81 | 97.04 | 98 | 97.45 | 97.71 | 97.53 |
| Rama-Z (Ramachandran plot Z-score, RMSD) | | | | | | | |
| whole | 0.25 (0.10) | -2.37 (0.14) | -0.32 (0.16) | -0.96 (0.16) | 0.07 (0.11) | 0.39 (0.15) | -0.39 (0.14) |
| helix | 0.79 (0.11) | -2.51 (0.11) | -0.43 (0.14) | -0.92 (0.14) | 0.41 (0.12) | 1.40 (0.18) | -0.11 (0.17) |
| sheet | 0.34 (0.14) | 0.53 (0.24) | 0.80 (0.26) | 0.67 (0.27) | 0.37 (0.16) | 0.12 (0.19) | 0.09 (0.21) |
| loop | -0.27 (0.11) | -1.24 (0.16) | -0.20 (0.19) | -0.66 (0.19) | -0.21 (0.11) | -0.19 (0.15) | -0.37 (0.14) |
| Rotamer outliers (%) | 0.99 | 0.63 | 1.43 | 1.33 | 1.38 | 1.3 | 1.24 |
| Cβ outliers (%) | 0.22 | 0.22 | 0.09 | 0.29 | 0.04 | NA | 0.11 |
| Peptide plane (%) | | | | | | | |
| Cis proline/general | 2.1/0.0 | 2.3/0.0 | 2.4/0.0 | 2.7/0.0 | 1.9/0.0 | 1.7/0.0 | 2.7/0.0 |
| Twisted proline/general | 0.0/0.0 | 0.0/0.0 | 0.0/0.0 | 0.0/0.0 | 0.0/0.0 | 0.0/0.0 | 0.0/0.0 |
| CaBLAM outliers (%) | 1.25 | 1.58 | 1.38 | 1.16 | 1.44 | 1.47 | 1.31 |
| ADP (B-factors) | | | | | | | |
| Iso/Aniso (#) | 150275/0 | 54113/0 | 52172/0 | 51204/0 | 142256/0 | 89703/0 | 87980/0 |
| min/max/mean | | | | | | | |
| Protein | 30.00/157.46/62.71 | 30.00/157.46/60.28 | 42.46/98.61/93.18 | 52.46/107.76/74.56 | 6.21/166.19/72.45 | 6.21/166.19/56.77 | 45.53/66.75/54.63 |
| Nucleotide | 44.82/72.30/59.60 | 34.29/68.86/53.18 | 13.92/341.26/85.44 | 50.28/102.49/69.38 | 13.92/545.91/88.78 | 22.67/545.91/90.50 | 45.86/64.80/53.21 |
| Ligand | 43.78/115.41/63.70 | 52.53/115.41/57.82 | 30.00/81.22/33.96 | 26.67/200.00/78.23 | 30.00/119.73/58.50 | 30.68/119.73/63.66 | 18.41/220.39/66.07 |
| Water | 39.19/109.28/63.66 | 49.85/109.28/52.63 | 30.00/117.87/47.57 | 0.65/60.94/39.21 | 30.00/117.87/53.08 | 30.00/66.94/50.30 | 0.65/89.73/36.00 |
| Occupancy | | | | | | | |
| Mean | 1 | 1 | 1 | 1 | 1 | 1 | 1 |
| occ = 1 (%) | 100 | 100 | 100 | 100 | 100 | 100 | 100 |
| 0 < occ < 1 (%) | 0 | 0 | 0 | 0 | 0 | 0 | 0 |
| occ > 1 (%) | 0 | 0 | 0 | 0 | 0 | 0 | 0 |
| Data | | | | | | | |
| Box | | | | | | | |
| Lengths (Å) | 230.26,247.68,268.96 | 194.15,243.16,160.6 | 143.1,226.84,202.46 | 142.04,226.84,205.64 | 268.18,269.24,234.26 | 182.32,228.96,234.26 | 182.32,227.9,233.2 |
| Angles (°) | 90,90,90 | 90,90,90 | 90,90,90 | 90,90,90 | 90,90,90 | 90,90,90 | 90,90,90 |
| Supplied Resolution (Å) | 2.2 | 2.1 | 2.9 | 3.1 | 2.5 | 2.2 | 2.2 |
| Resolution Estimates (Å) | Masked, Unmasked | Masked, Unmasked | Masked, Unmasked | Masked, Unmasked | Masked, Unmasked | Masked, Unmasked | Masked, Unmasked |
| d FSC (half maps; 0.143) | ---,--- | ---,--- | --,-- | --,-- | --,-- | --,--- | --,-- |
| d 99 (full/half1/half2) | 2.9/---/---, 2.8/---/--- | 3.0/---/---, 2.9/---/--- | 3.1/--/--, 2.9/--/-- | 3.6/--/--, 3.4/--/-- | 3.1/--/--, 3.0/--/-- | 2.9/--/--, 2.7/--/-- | 2.9/---/---, 2.7/---/--- |
| d model | 2.6, 2.6 | 2.7, 2.7 | 2.5, 2.8 | 3.1, 3.1 | 2.7, 2.7 | 2.7, 2.6 | 2.6, 2.6 |
| d FSC model (0/0.143/0.5) | 2.1/2.2/2.6, 2.2/2.2/2.8 | 2.1/2.1/2.4, 2.1/2.2/2.9 | 2.5/3.0/4.2, 2.8/3.3/6.8 | 3.0/3.0/3.4, 3.0/3.1/4.1 | 2.5/2.5/2.9, 2.5/2.6/3.1 | 2.2/2.2/2.6, 2.2/2.2/2.8 | 2.1/2.2/2.6, 2.2/2.2/2.8 |
| Map min/max/mean | -0.10/0.38/0.01 | -0.08/0.30/0.01 | -0.13/0.44/0.02 | -0.08/0.26/0.01 | -0.13/0.47/0.01 | -0.16/0.55/0.01 | -0.18/0.54/0.01 |
| Model vs. Data | | | | | | | |
| CC (mask) | 0.8 | 0.85 | 0.66 | 0.83 | 0.79 | 0.85 | 0.84 |
| CC (box) | 0.84 | 0.65 | 0.63 | 0.74 | 0.83 | 0.84 | 0.84 |
| CC (peaks) | 0.75 | 0.42 | 0.32 | 0.64 | 0.76 | 0.79 | 0.79 |
| CC (volume) | 0.8 | 0.85 | 0.67 | 0.83 | 0.8 | 0.85 | 0.84 |
| Mean CC for ligands | 0.72 | 0.69 | 0.4 | 0.78 | 0.6 | 0.75 | 0.7 |

**Cryo-EM reveals multiple mechanisms of ribosome inhibition by doxycycline**

**Stuart et al.**

**Supplementary Figure 1: Electron density demonstrates a novel doxycycline binding site in *C. burnetii*.**

(**a,b**) Following classification, clear density could be observed for doxycycline in the A-site of the ribosomal 30S subunit of *C. burnetii* and *E. coli* respectively (rRNA and doxycycline shown as sticks. Colours: oxygen, red; nitrogen, blue; phosphorus, orange; rRNA carbon, cyan; doxycycline carbon, green; electron density, grey). The coordinating rRNA base C1054 is highlighted in both structures. O11 and O12 of doxycycline coordinate a magnesium bound by the phosphate groups of C1054 and G1198 while ring D stacks upon the base of C1054. Another magnesium bound to the phosphate of A966 coordinates O3. (**c**) A second doxycycline binding site is observed in the *C. burnetii* 50S ribosome. Three doxycycline molecules form a stack with three coordinated magnesium atoms. Electron density shown as a blue net. (**d**) The respective *E. coli* region following classification shows little evidence of an equivalent state, with limited evidence for only the lower molecule in the image shown. Colours: doxycycline carbon, white; electron density, grey net.

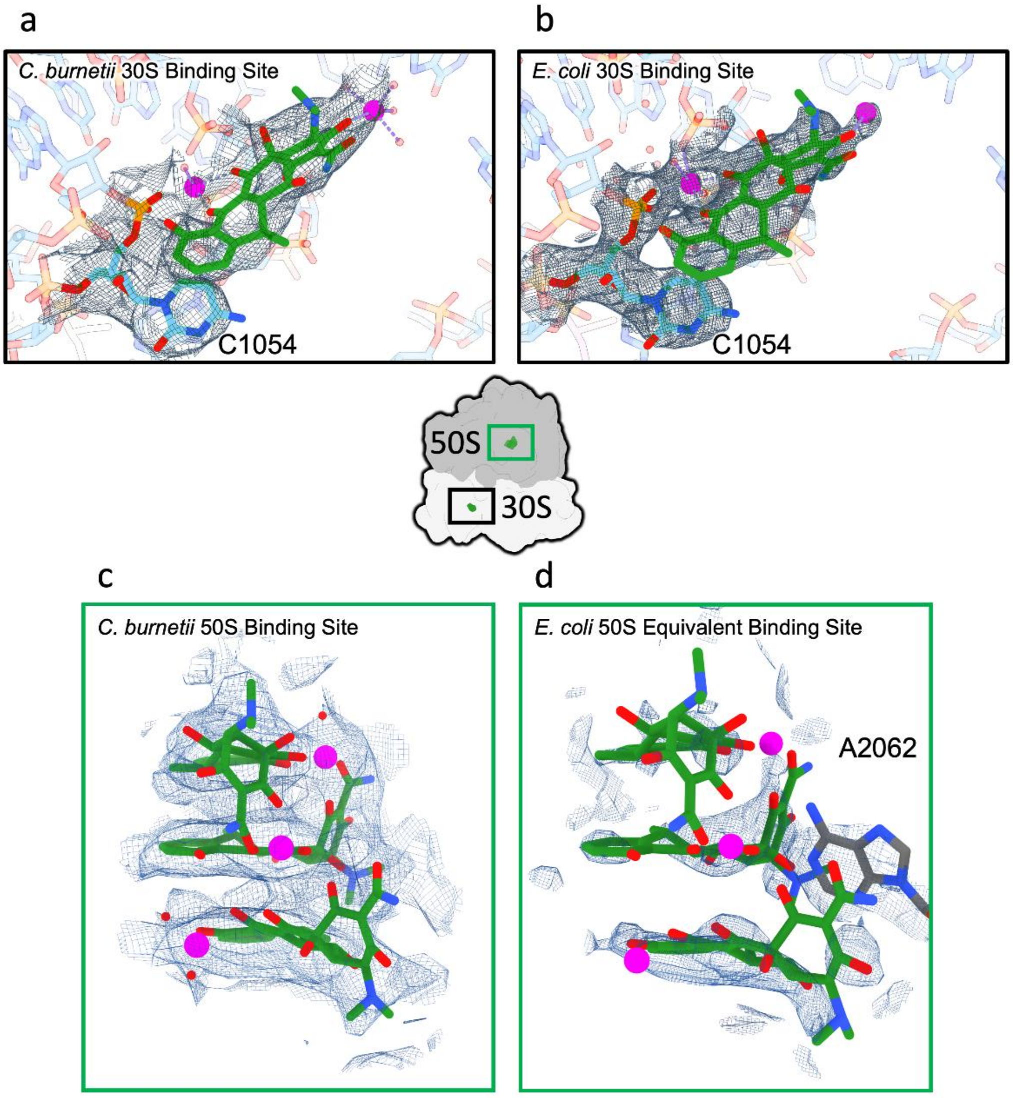

**Cryo-EM reveals multiple mechanisms of ribosome inhibition by doxycycline**

**Stuart et al.**

**Supplementary Figure 2: rRNA modifications observed in the *C. burnetii* ribosome.**

*C. burnetii* shows fewer rRNA post-translational modifications than *E. coli*. Modifications were included in the structure where they were clearly indicated by the density. Modifications were only included if an orthologue of an *E. coli* modification gene is present in *C. burnetii* and supported by map density (e.g. pseudo uridylation have been excluded, even if a corresponding *C. burnetii* gene is present). Modifications are numbered in the *C. burnetii* ribosome, with *E. coli* numbering in parentheses. Modification abbreviations: OMU: 2'-*O*-methyluridine; 2MA: 2-methyladenosine; 7MG: 7-methylguanosine; 5MC, 5-methylcytosine; 4,2MC: 4-*N*-methyl, 2'-*O*-methylcytosine; 3MU, 3-*N*-methyluridine; 2MG, 2-*N*-methylguanosine; 6M (2)A, 6-*N*,*N*-dimethyladenine. Bases shown as sticks. Colours: yellow, carbon; red, oxygen; blue, nitrogen; light blue, *E. coli* density; grey, *C. burnetii* density.

| *C. burnetii* Modification (*E. coli*) | *E. coli* | *C. burnetii* | *C. burnetii* Gene | |
|---|---|---|---|---|
| OMU-2572 (2552) | 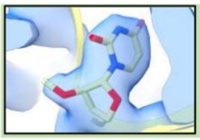 | 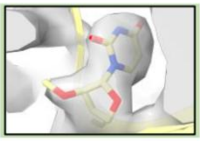 | *rlmE, CBU_1353* | **23S** |
| 2MA-2523 (2503) | 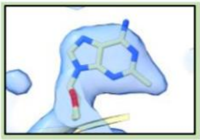 | 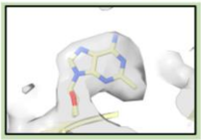 | *rlmN, CBU_1252* | |
| OMG-2271 (2251) | 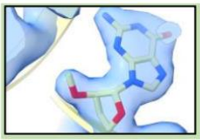 | 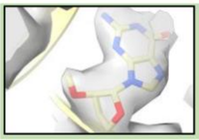 | *rlmB, CBU_0986* | |
| 7MG-524 (527) | 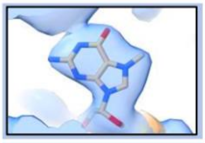 | 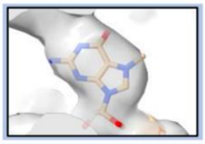 | *rsmG, CBU_1925* | **16S** |
| 2MG-962 (966) | 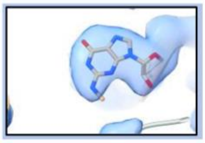 | 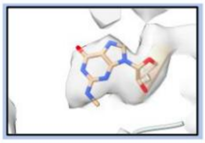 | *rsmD, CBU_1899* | |
| 5MC-963 (967) | 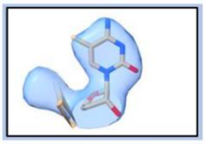 | 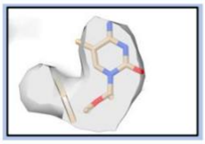 | *rsmB, CBU_1915* | |
| 4,2MC-1399 (1402) | 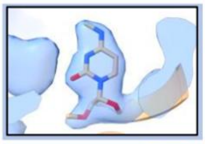 | 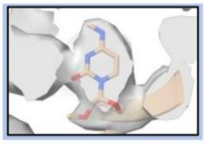 | *rsmH & rsmI, CBU_0116 & CBU_1739* | |
| 3MU-1495 (1498) | 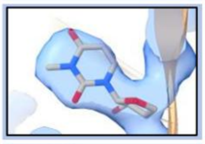 | 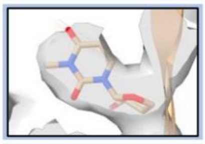 | *rsmE, CBU_1960* | |
| 2MG-1513 (1516) | 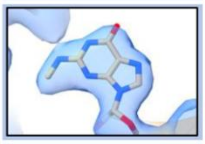 | 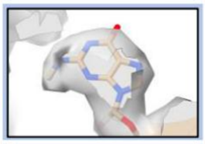 | *rsmJ, CBU_1876* | |
| 6M(2)A-1515 (1518) | 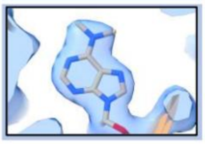 | 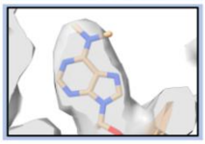 | *rsmA, CBU_1982* | |

**Cryo-EM reveals multiple mechanisms of ribosome inhibition by doxycycline**

**Stuart et al.**

**Supplementary Figure 3: HPF$_{cold}$ binds the 30S subunit of the ribosome, protecting the anti-Shine Dalgarno sequence.**

Structural characterisation of HPF$_{cold}$ bound to the *C. burnetii* ribosome. (**a**) The cold shock domain (CSD) interacts intimately with the 3' region of the 16S rRNA containing the anti-Shine Dalgarno (aSD) sequence at the mRNA entrance channel. rRNA shown as ribbon and sticks with the aSD sequence in rainbow colours; CSD shown as cartoon and semi-transparent surface in blue. (**b**) The aSD wraps around the CSD, with U1534 stacking upon Y142 while C1535 stacks with F132 and forms a hydrogen bond pseudo-pair to R123 with its N3 and O2 atoms. rRNA and interacting residues of CSD shown as sticks. Colours: oxygen, red; nitrogen/CSD carbon, blue; phosphate/rRNA carbon, orange. (**c**) R177 acts as a bridge between two rRNA segments, forming salt bridges with a cleft of the rRNA phosphate backbone formed by G924-925 and C1530. rRNA shown as ribbon and sticks (carbons and ribbon in cyan). aSD not differentiated with colour but C1535 indicated with open black arrowhead. (**d**) HPF$_{cold}$ and bS21 pincer the aSD. Further interactions are made on the other face of the CSD with rRNA (yellow) and bS7. Ribosomal proteins shown as pink cartoon.

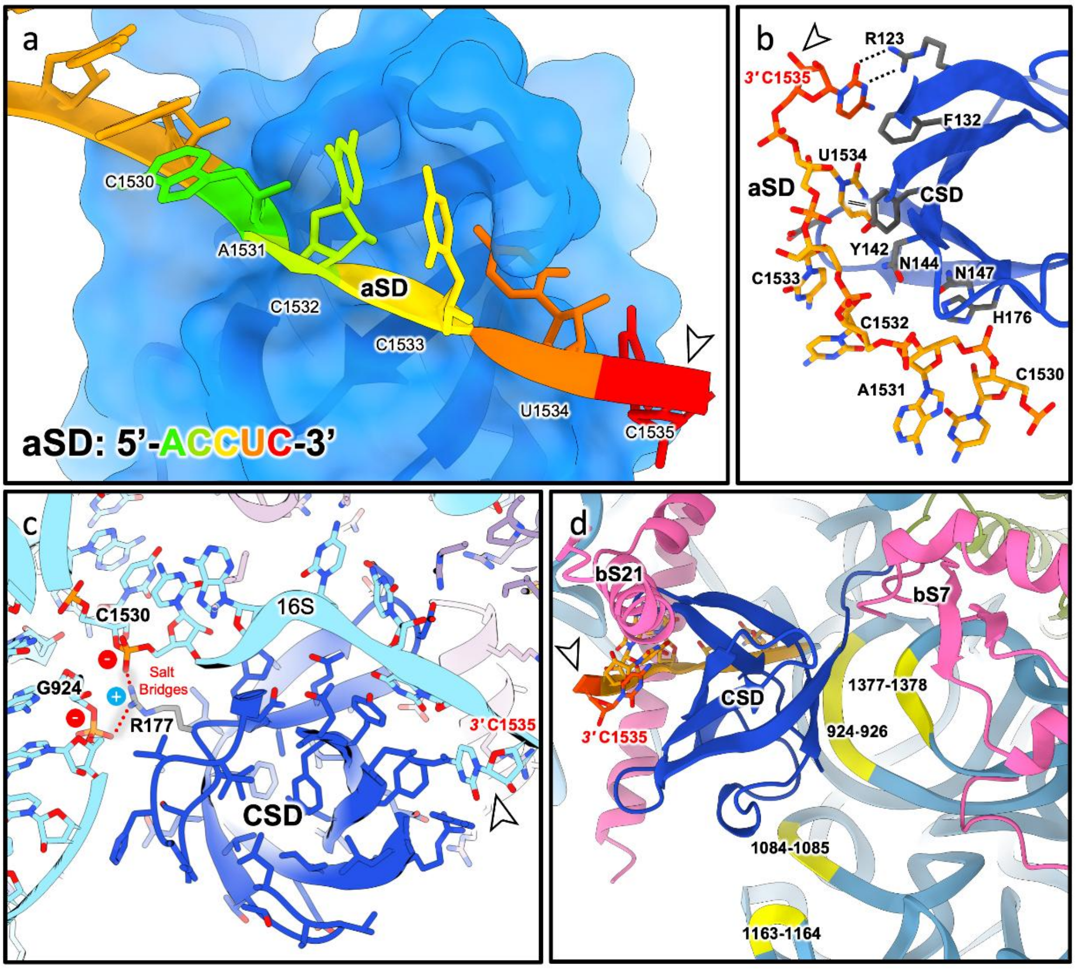

**Cryo-EM reveals multiple mechanisms of ribosome inhibition by doxycycline**

**Stuart et al.**

**Supplementary Figure 4: $HPF_{cold}$ is an additional hibernation promoting factor**

While no structural data for sequence homologues could be identified in the protein data bank, the CSD displays a classical cold shock protein fold. (**a**) Sequence alignment (Figure S5a) against the *E. coli* (K12) cold shock proteins reveals generally low sequence identity, indicating evolutionary divergence. (**b**) AlphaFold 3 [1] predicts that *Legionella pneumophila* $HPF_{cold}$ is structurally highly comparable to $CbHPF_{cold}$. Proteins shown as cartoon; green, *C. burnetii*, blue, *L. pneumophila*. (**c**) A KLEVG motif (magenta) is conserved between *C. burnetii* and *L. pneumophila* $HPF_{cold}$ CSDs (blue) and not present in standalone cold shock proteins. This motif falls on the CSD interface contacting helix 40 of the 16S rRNA, opposite to the aSD interface (orange). The KLEVG Lys155 inserts into the backbone cleft between A1164 and G1084. Proteins and rRNA shown as cartoon with semi-transparent surface, with sticks for the KLEVG motif. (**d**) Structural alignment between $HPF_{cold}$ CSD and CspA from *E. coli* (PDB: pdb_00001mjc [2]). RMSD values (calculated in ChimeraX 1.9 [3]) sequence alignment score = 107.2, RMSD between 56 pruned residues is 0.799 Å; and 1.756 Å across all 66 aligned residues. (**e**) Structural comparison of known ribosomal hibernation factors (alignment: Figure S5b). Structures are superposed on the common domain and shown as cartoon. Colours: $HPF_{short}$, orange (PDB: pdb_00004v8i [4]); YfiA, blue (PDB: pdb_00004v8h [4]); $HPF_{long}$, magenta (PDB: pdb_00005myj [5]); $HPF_{cold}$, green and blue. Both the N-terminal (NTD) and C-terminal (CTD) domains of $HPF_{long}$ are indicated. Both $HPF_{cold}$ and $HPF_{long}$ C-terminal domains likely function to preserve the ribosome during hibernation, albeit in 70S and 100S ribosomal forms respectively.

a

| *E.coli* (K12 Csp) | Max Score | Total Score | Query Cover | E value | Per. Ident | Acc. Len |
|---|---|---|---|---|---|---|
| CspA | 25.0 | 25.0 | 88% | 1e-05 | 31% | 70 |
| CspB | 20.0 | 20.0 | 85% | 0.001 | 28% | 71 |
| CspC | 24.6 | 24.6 | 94% | 2e-05 | 29% | 69 |
| CspD | 22.7 | 22.7 | 88% | 1e-04 | 26% | 74 |
| CspE | 23.1 | 23.1 | 94% | 6e-05 | 29% | 69 |
| CspF | - | - | - | - | - | - |
| CspG | 19.6 | 19.6 | 85% | 0.001 | 30% | 70 |
| CspH | 16.9 | 16.9 | 85% | 0.013 | 32% | 70 |
| CspI | 20.8 | 20.8 | 34% | 5e-04 | 46% | 70 |

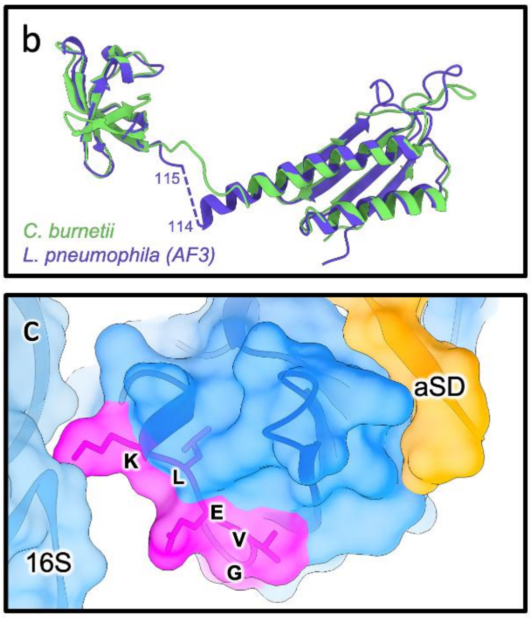


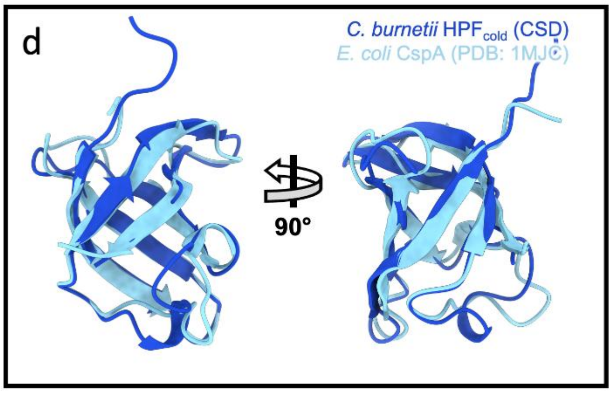


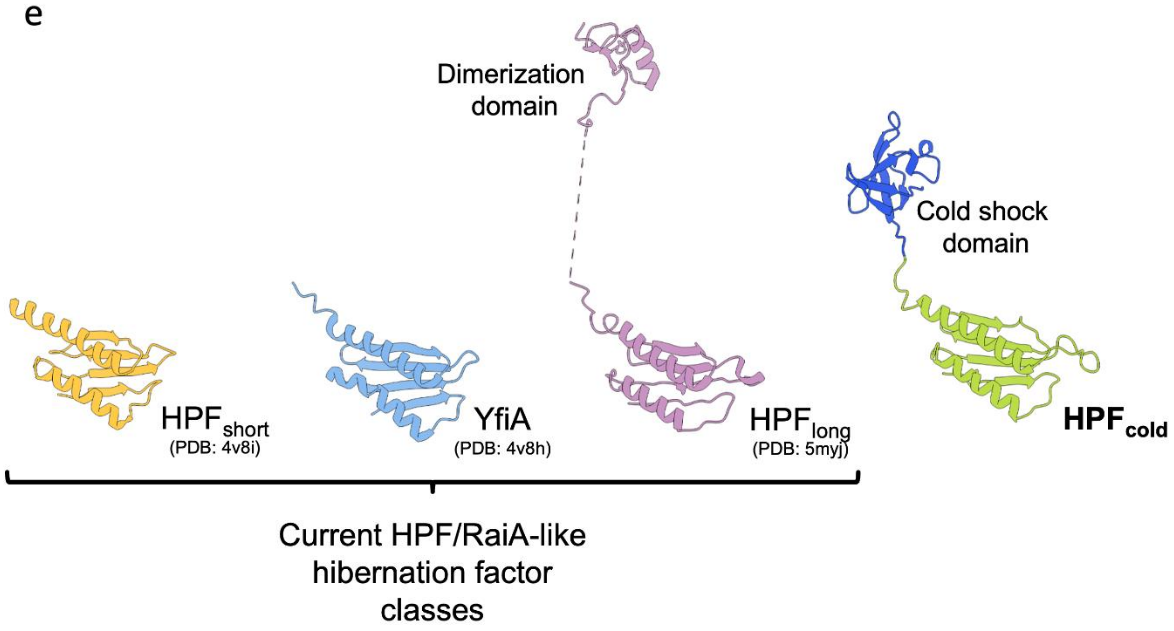

**Cryo-EM reveals multiple mechanisms of ribosome inhibition by doxycycline**

**Stuart et al.**

**Supplementary Figure 5: Alignment of $HPF_{cold}$ with paralogues of the two domains**

(**a**) The cold shock domain (CSD, Glu115 onwards) of $HPF_{cold}$ from *Coxiella burnetii* and *Legionella pneumophila* was aligned to the nine cold shock proteins of *E. coli* K12 (Figure 4a). (**b**) The hibernation factor domain of $HPF_{cold}$ from *C. burnetii* and *L. pneumophila* was aligned to example sequences of the three established hibernation factors ($HPF_{long}$ from *Staphylococcus aureus*, YfiA from *E. coli* K12, and HPFshort from *E. coli* K12. Alignments were performed using Clustal Omega [6] and visualised using Espript 3 [7].

**Cryo-EM reveals multiple mechanisms of ribosome inhibition by doxycycline**

**Stuart et al.**

a

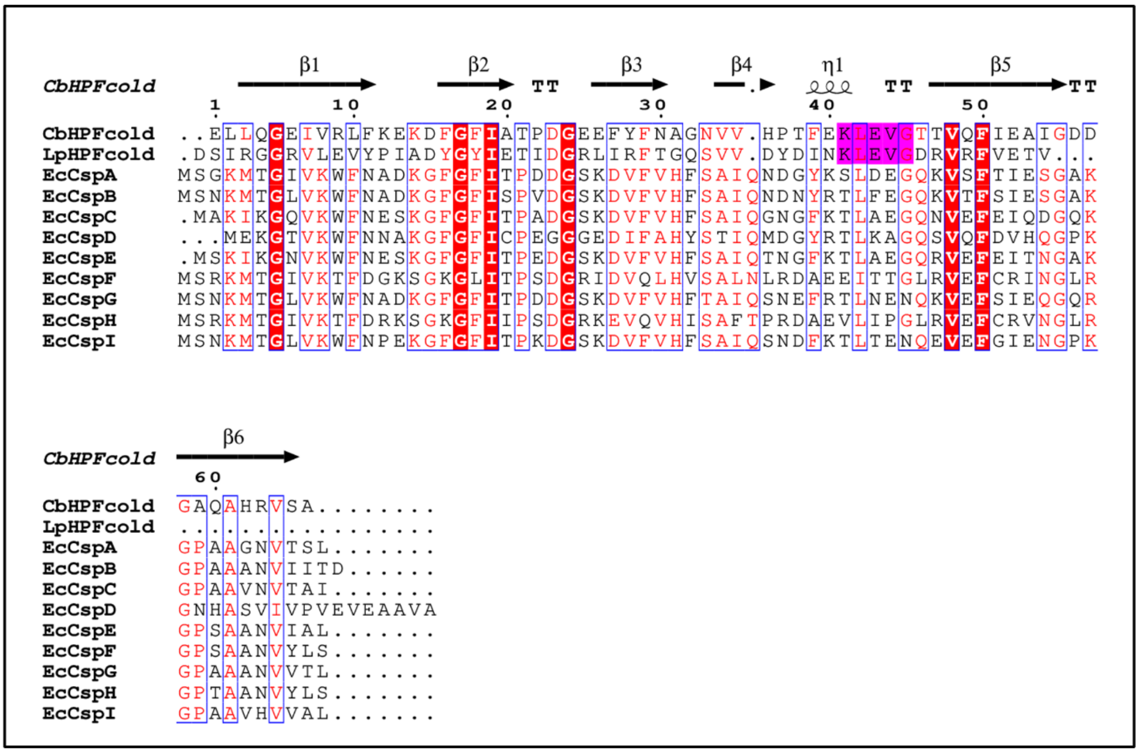


b

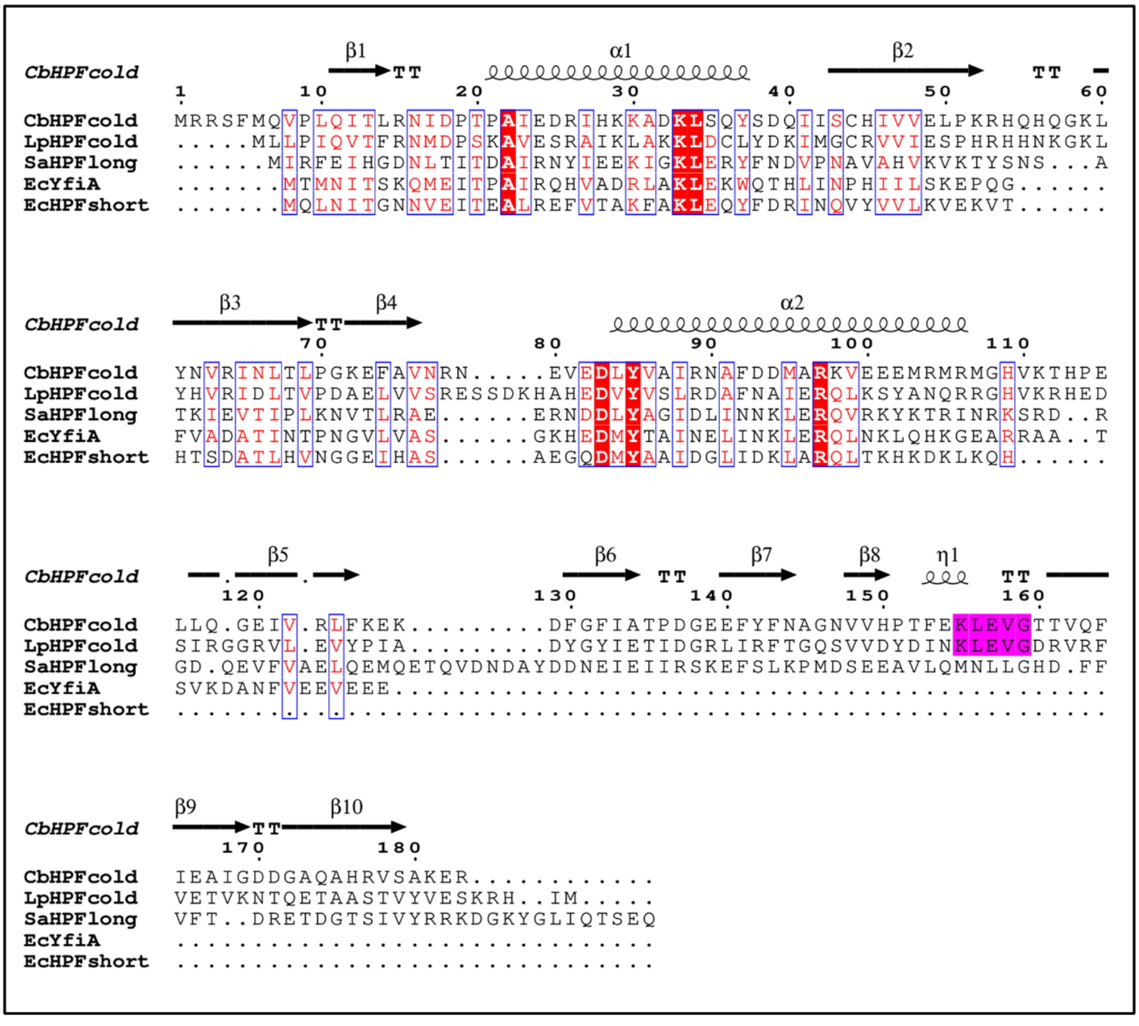

**Cryo-EM reveals multiple mechanisms of ribosome inhibition by doxycycline**

**Stuart et al.**

**Supplementary Figure 6: CLaSP binding in context**

(**a**) *Coxiella burnetii* displays a truncated 5S rRNA in comparison to *Escherichia coli*, more similar to the 5S found in *Mycobacterium tuberculosis*. Like *C. burnetii*, *M. tuberculosis* also occupies this cleft with a ribosomal protein, named bL37 (pdb_00005xym). (**b**) bL37 displays a more compact, helical structure than CLaSP. 5S rRNA shown as purple ribbon and sticks with semi-transparent surface. 23S rRNA shown as light green ribbon. CLaSP (pink) and bL37 (green) shown as cartoon and sticks with semi-transparent surface. Other ribosomal proteins shown as grey cartoon and semi-transparent surface. The C-terminal residue (A24) of bL37 is at the rear of the protein as shown (black arrow). (**c**) CLaSP is located adjacent to *nusB*, one base out of frame from the 3' end. Orange – *nusB*; pink – *CLaSP*.

**Cryo-EM reveals multiple mechanisms of ribosome inhibition by doxycycline**

**Stuart et al.**

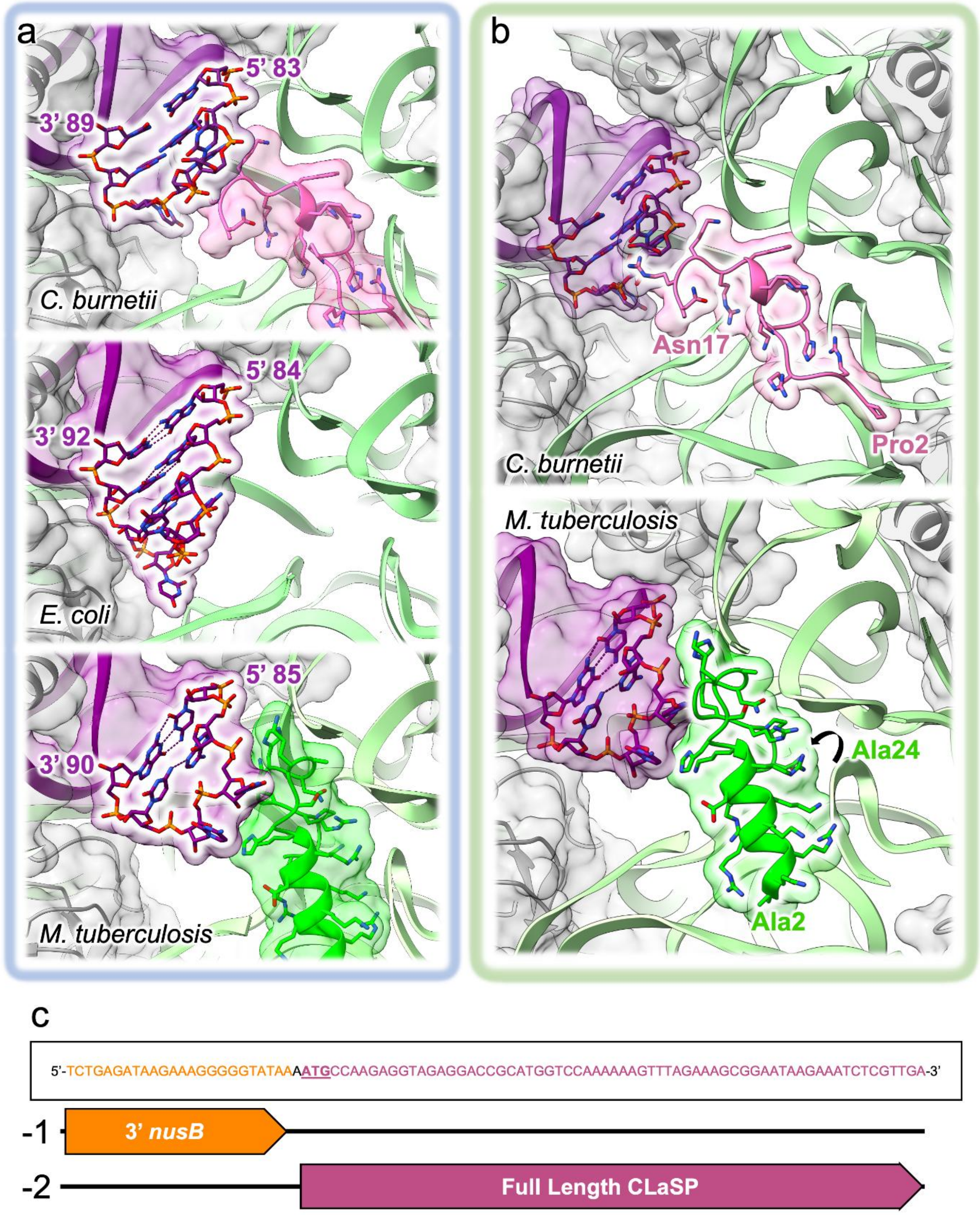

**Cryo-EM reveals multiple mechanisms of ribosome inhibition by doxycycline**

**Stuart et al.**

**Supplementary Figure 7: *C. burnetii* data processing pipeline**

Overview of the data processing strategy. Images comparing electron density following classification are at the same sigma values. Left: a mask was applied at the site of the doxycycline triple stack (purple area shown). This separated the particles into five classes, with two classes (41.5% of particles) representing the doxycycline bound 50S ribosome and 58.5% representing an empty 50S ribosome. Right: a mask was applied at the site of $HPF_{cold}$ (cyan area shown). Two classes were separated, with 55% bound to $HPF_{cold}$ (purple) and 45% showing doxycycline (green) and a tRNA (blue).

**Cryo-EM reveals multiple mechanisms of ribosome inhibition by doxycycline**

**Stuart et al.**

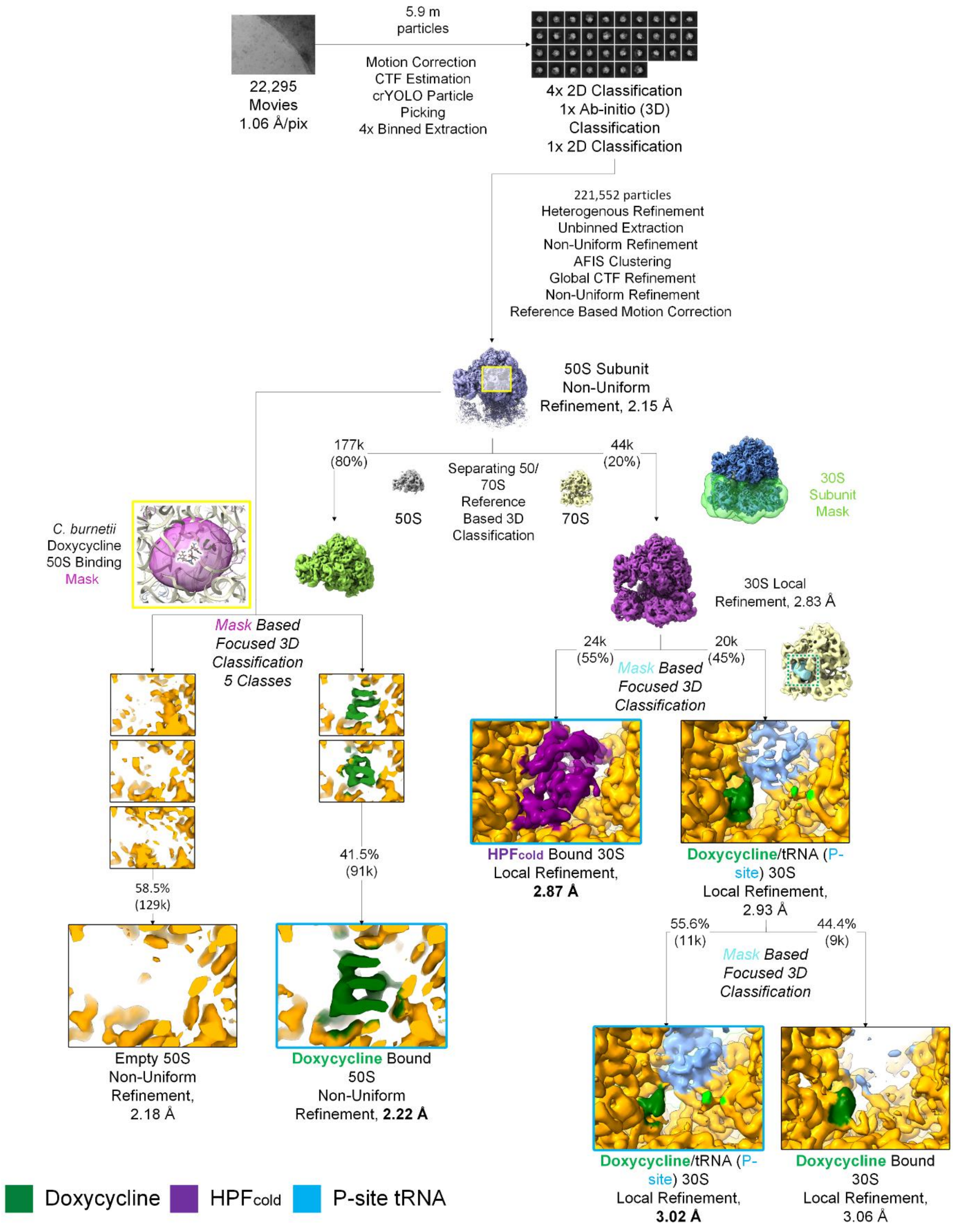

**Cryo-EM reveals multiple mechanisms of ribosome inhibition by doxycycline**

**Stuart et al.**

**Supplementary Figure 8: *E. coli* data processing pipeline**

Overview of the data processing strategy. Images comparing electron density following classification are at the same sigma values. Left: a mask was applied at the site of the doxycycline binding to the 50S ribosome (purple area shown). This separated the particles into ten classes, with one class (12.3% of particles) representing the doxycycline bound 50S ribosome. Right: a mask was applied at the site of YfiA (hibernation factor) (cyan area shown). Two classes were separated, with 61% bound to YfiA (purple) and 39% showing doxycycline (green) and a tRNA(blue).

**Cryo-EM reveals multiple mechanisms of ribosome inhibition by doxycycline**

**Stuart et al.**

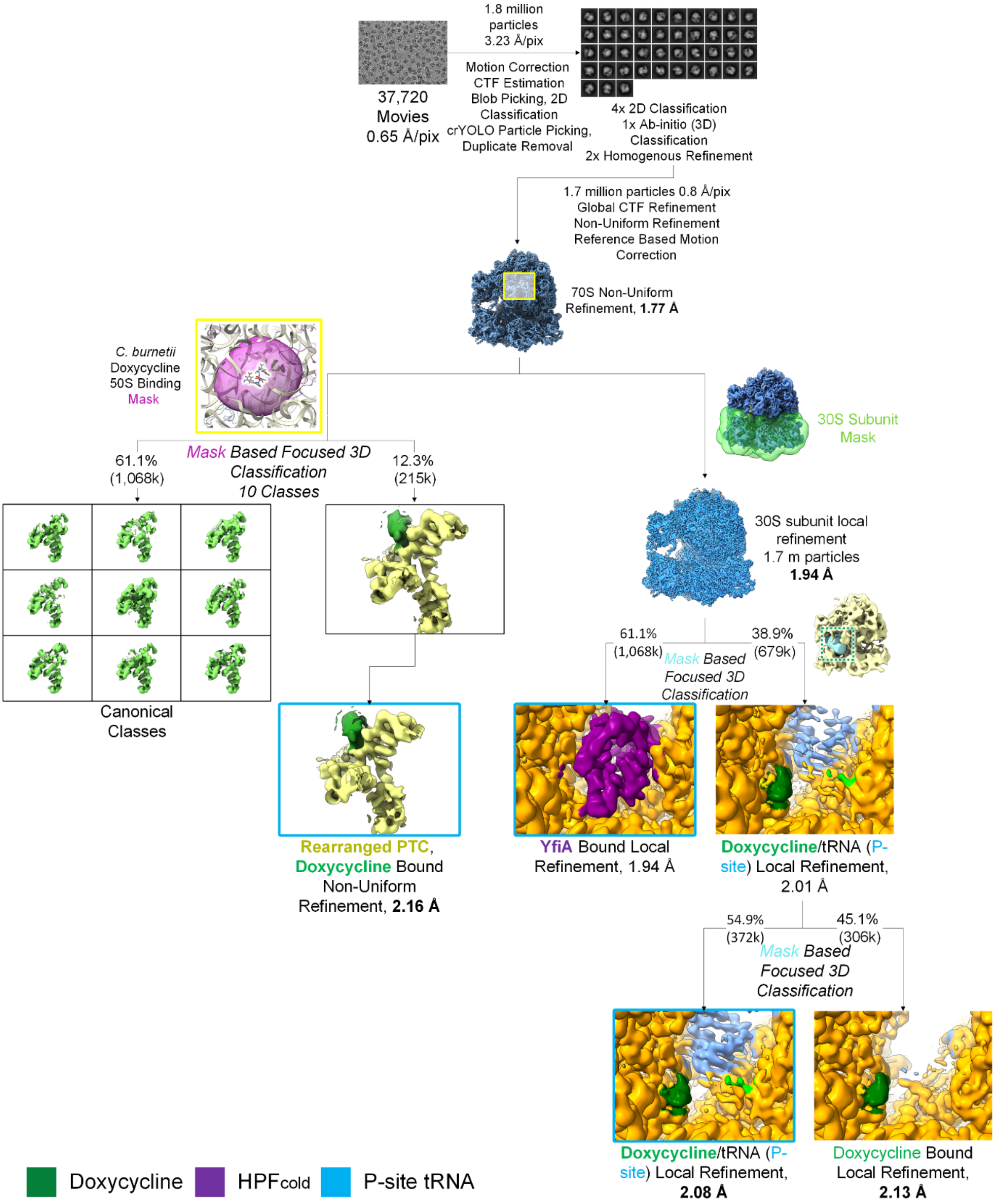

**Cryo-EM reveals multiple mechanisms of ribosome inhibition by doxycycline**

**Stuart et al.**

**Supplementary Figure 9: Electron density for doxycycline bound in the exit tunnel, comparison to the empty exit tunnel**

(**a, b**) Focused classification of the *Coxiella burnetii* 50S ribosome (Figure S7) separated the particles into those containing a three-molecule doxycycline stack (**a**) and doxycycline-free 50S ribosomes (**b**) reconstructions. (**c**) A striking novel interaction is between DOX2 and U2526, stabilising the conformation of the base in a luminal, translationally stalled-like position. (**d**) Equivalent density for U2526 in the unbound ribosome is weak, suggesting that U2526 samples multiple conformations. (**e**) Aligning the macrolide resistance conferring m2m8A2503 modified ribosome reveals that the C8 methylation is distant from the antibiotic binding (**c**). This suggests that the m2m8A2503modification may not confer resistance to this doxycycline binding mode. (**f,g**) The doxycycline bound (**f**) and unbound (**g**) exit tunnels additionally differ through an absence in waters coordinated to A2077 and A2078 in the bound tunnel, which are readily observed in the unbound exit tunnel (open arrowheads). These may be displaced upon antibiotic binding. 23S rRNA and doxycycline shown as sticks; electron density shown as blue mesh. Colours: nitrogen, blue; oxygen, red; phosphorus, orange; rRNA carbon, light green; doxycycline carbon, dark green.

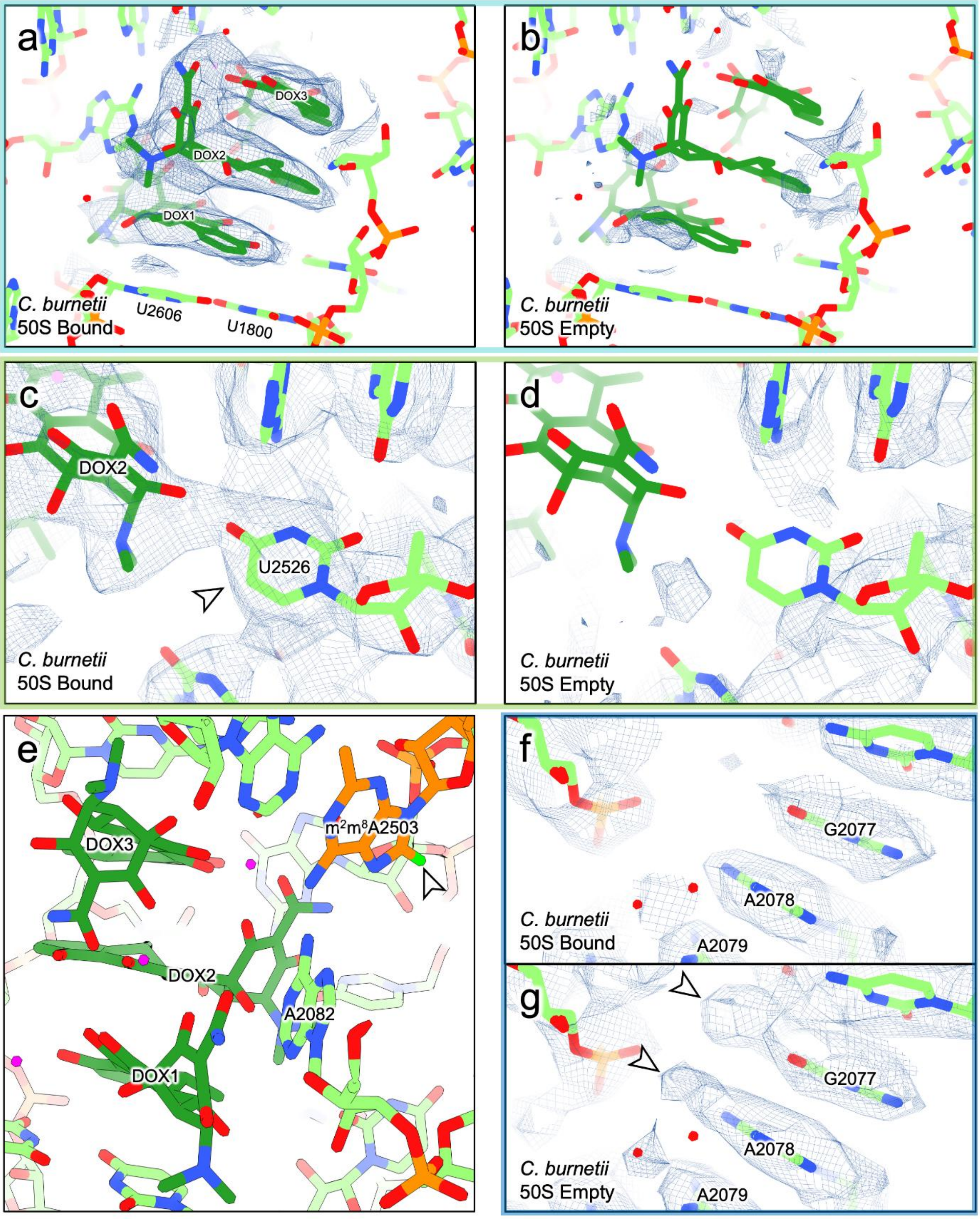
a
DOX3
DOX2
DOX1
C. burnetii
50S Bound
U2606
U1800
b
C. burnetii
50S Empty
c
DOX2
U2526
C. burnetii
50S Bound
d
C. burnetii
50S Empty
e
DOX3
m2m8A2503
DOX2
A2082
DOX1
f
G2077
A2078
C. burnetii
50S Bound
A2079
g
G2077
A2078
C. burnetii
50S Empty
A2079

**Cryo-EM reveals multiple mechanisms of ribosome inhibition by doxycycline**

**Stuart et al.**

**Supplementary Figure 10: Structural details of doxycycline within the *C. burnetii* exit tunnel**

(**a**) Doxycycline stacking is reliant upon three magnesium ions. The "top" ion (dashed red box, top left) is coordinated by two oxygen atoms each from DOX2 and DOX3 and two water molecules; the central magnesium ion (blue dashed box, bottom left) is coordinated by all three molecules, with each doxycycline donating two oxygen atoms to complete the octahedral magnesium coordination. The third magnesium is coordinated only by DOX1 (bottom right). DOX1 shows a strong π-interaction with A2082 (green dashed box, top right). 23S rRNA and doxycycline shown as sticks; magnesium and water shown as spheres. Colours: nitrogen, blue; oxygen, red; phosphorus, orange; rRNA carbon, light green; doxycycline carbon, dark green; magnesium, pink.
(**b**) Modelling the triple stacking with later generation tetracyclines indicates the stacking we observe for doxycycline is unlikely to be feasible due to clashes with the walls of the exit tunnel. (**c-e**) Previous investigations have displayed tetracyclines and tetracycline-like molecule binding somewhat in similar locations to DOX1, such as sarecycline and tetracenomycin-X. Sarecycline bound to the *Cutibacterium acnes* ribosome (**c**; pdb_00008cvm [8]) shows the 23S backbone encroaching into the doxycycline stacking space (arrowhead), with sarecycline in a different pose to DOX1 (**d**). Tetracenomycin X bound to the *E. coli* ribosome (**e**; pdb_00006y69 [9]) binds in a similar location to DOX1 but in a very different pose that would not support the magnesium coordination that characterises the doxycycline triple stack. Antibiotics and selected rRNA bases shown as sticks, rRNA as ribbon; sarecycline carbons, salmon; *C. acnes* rRNA, blue; tetracenomycin X, light blue; *E. coli* rRNA, cyan. (**f,g**) Doxycycline recapitulates the hydrogen bond to A2078 (A2058 in *E. coli*) that characterises macrolide binding to the ribosome exit tunnel (**f**; pdb_00006xhx [10]). Here, a water coordinated by the DOX2-DOX3 magnesium is taking the role of the hydroxyl group of the macrolide (**g**). Erythromycin and 23S rRNA shown as sticks; erythromycin carbons, grey; *E. coli* rRNA, light orange.

**Cryo-EM reveals multiple mechanisms of ribosome inhibition by doxycycline**

**Stuart et al.**

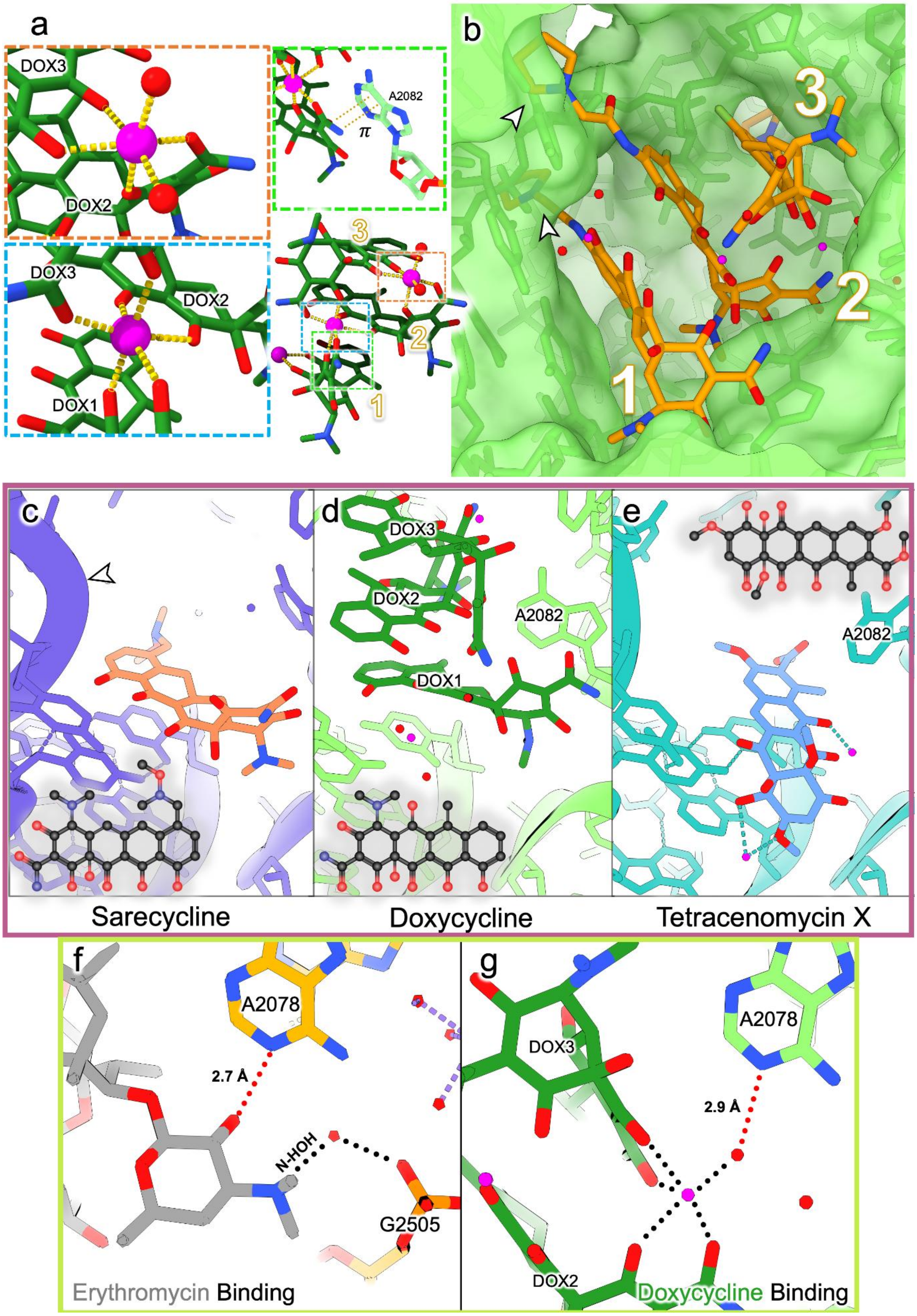

**Cryo-EM reveals multiple mechanisms of ribosome inhibition by doxycycline**

**Stuart et al.**

**Supplementary Figure 11: Inactive *E. coli* ribosome identified through doxycycline driven serendipity**

In depth classification of our *E. coli* data to identify *C. burnetii*-like binding revealed a subset displaying a substantially rearranged, inactive ribosome. (**a**) Comparison of the unbound (upper) and doxycycline bound (lower) *E. coli* ribosome conformations, showing only nucleotides that display motion in response to doxycycline binding. rRNA shown as ribbon with riboses and bases as sticks. Doxycycline shown as sticks with semi-transparent green surface. Colours: rRNA carbon and ribbon, orange; doxycycline carbon, green; oxygen, red; nitrogen, blue. (**b**) The inactive conformation is incompatible with the typical bL4extended loop structure (upper). In the doxycycline bound ribosomes this region is unstructured (lower; open arrowhead). (**c**) tRNA blocking nucleotides are stabilised by new stacking interactions. U2585 stacks upon C2063 and U2506 stacks upon A2602. To achieve this, both undergo a rotation from their canonical tRNA bound conformation (blue dotted outline) upon doxycycline binding (yellow dotted outline). Green arrows indicate movement of U2585 and A2602 while yellow arrows indicate movement of C2063 and U2506. Translucent neighbouring bases show that doxycycline conformational change is limited only to U2585 and A2602.

The doxycycline bound population had only a minor contribution (~12%, figure S8) to the bulk 1.77 Å reconstruction, with occasional weak density which would have been ignored were it not for the particular attention paid to this area, driven by the *C. burnetii* ribosome. This is a specific strength of cryo-EM as a technique and demonstrates that there may be more biological phenomena hidden in data already collected and deposited.

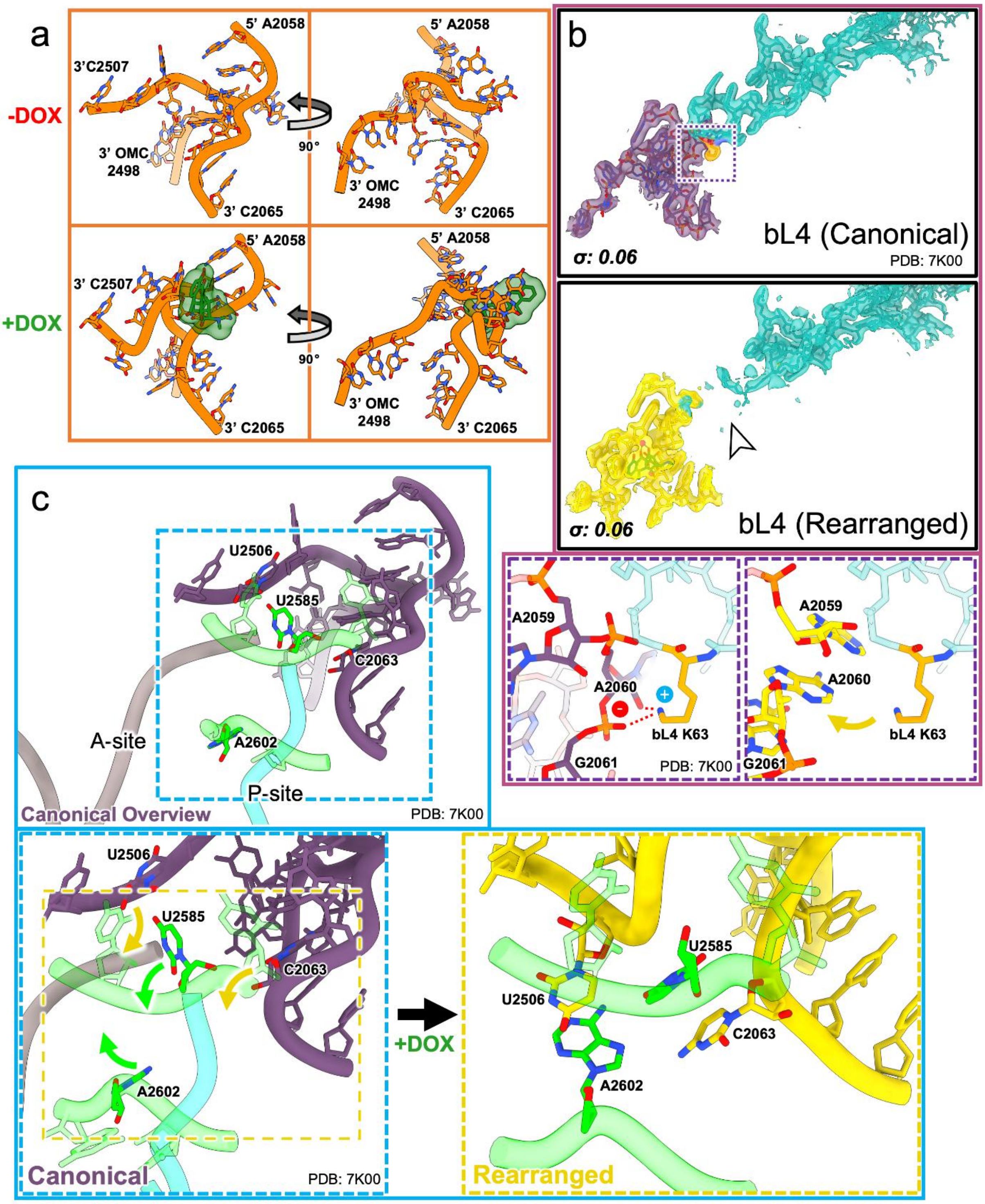
a
-DOX
+DOX
5' A2058
3'C2507
3' OMC 2498
3' C2065
90°
3' C2507
b
bL4 (Canonical)
σ: 0.06
PDB: 7K00
bL4 (Rearranged)
A2059
A2060
G2061
bL4 K63
c
U2506
U2585
C2063
A2602
A-site
P-site
Canonical Overview
Canonical
+DOX
Rearranged

**Cryo-EM reveals multiple mechanisms of ribosome inhibition by doxycycline**

**Stuart et al.**

**Supplementary Figure 12: Novel rRNA interactions are made in the doxycycline bound *E. coli* ribosome**

(**a,b**) Detailed view of the intra-rRNA interactions made. Only nucleotides that show rearrangement on doxycycline binding are shown (as in Figure S11a). G2502 and G2505 form a new stacking interaction (open arrowheads; highlighted in panel **c**). G2502 moves almost 30 Å (28.4 Å) when compared to the canonical ribosome (measuring N1-N1). Similarly, A2059, A2060, G2061 and C2501 are also forming a series of new stacking interactions (black arrowhead; highlighted in panels **d** and **e**). 2MA2503 forms a new stacking interaction with A2058 (grey arrowhead; highlighted in **f**). With C2063 flipped outwards towards incoming tRNAs, there is a cascade of new stacking interactions between A2062, C2064 and C2065 (black arrows). rRNA shown as ribbon with bases and riboses as sticks. Colours: carbon and ribbon, yellow; oxygen, red; nitrogen, blue. 2MA: 2-methyladenosine; y, pseudouridine; OMC, 2'-*O*-methylcytosine.

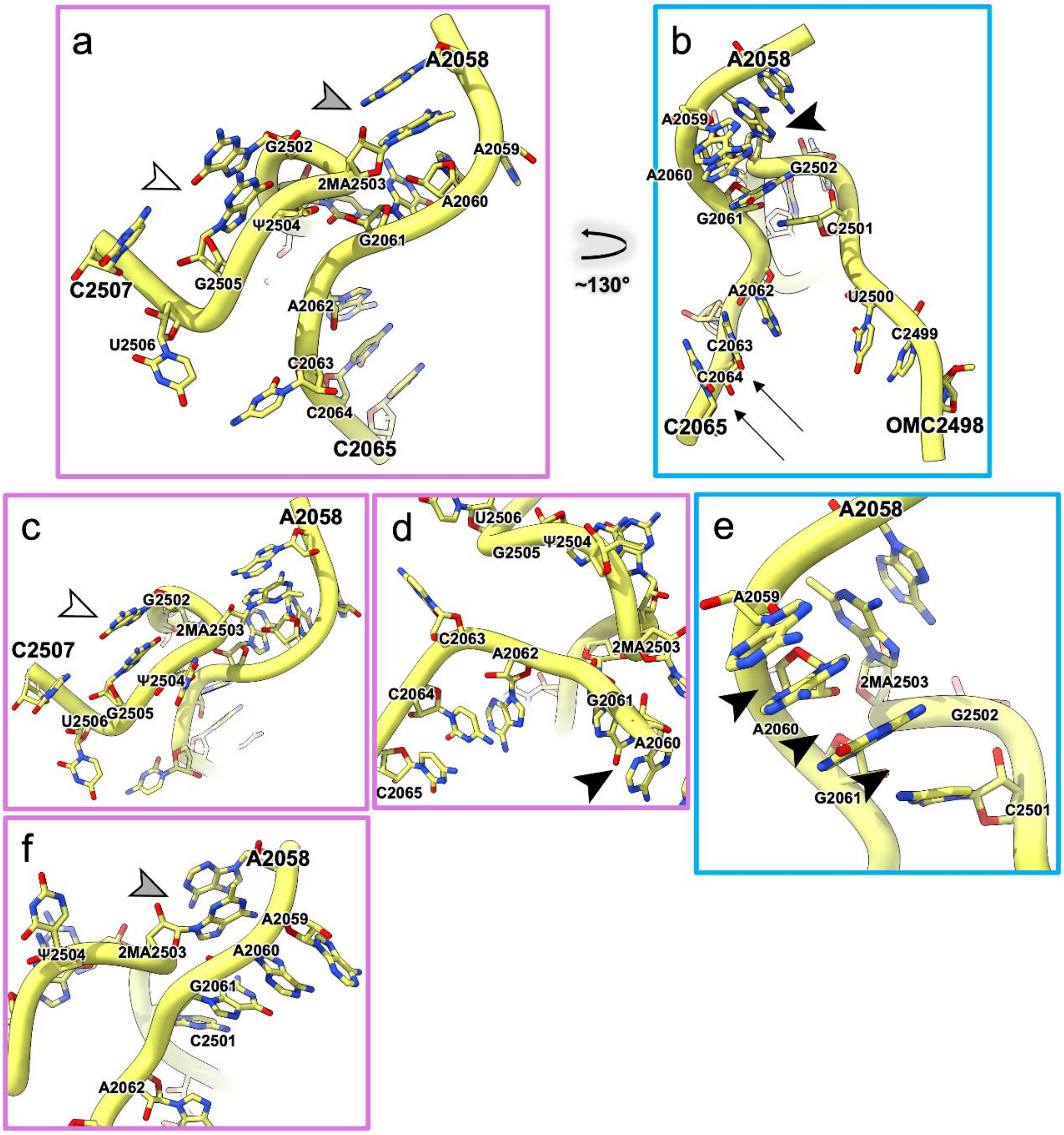

**Cryo-EM reveals multiple mechanisms of ribosome inhibition by doxycycline**

**Stuart et al.**

**Supplementary Figure 13: Cryo-EM map and model validation**

Gold standard Fourier shell correlation (FSC) curves for key refined maps from cryoSPARC [11]. Green box corresponds to *Coxiella burnetii* data (**a-e**) and turquoise to *Escherichia coli* (**f-g**). All resolutions called using the 0.143 FSC cut-off.

(**a**) Non-uniform refinement of the doxycycline bound 50S subunit. (**b**) Local refinement of the HPFcold bound 30S subunit. (**c**) Local refinement of the doxycycline bound 30S subunit. (**d**) Non-uniform refinement of the apo 50S subunit. (**g**) Non-uniform refinement of the 70S subunit. (**f**) Non-uniform refinement of the doxycycline bound rearranged *E. coli* ribosome. (**g**) Local refinement of the doxycycline bound 30S subunit.

**Cryo-EM reveals multiple mechanisms of ribosome inhibition by doxycycline**

**Stuart et al.**

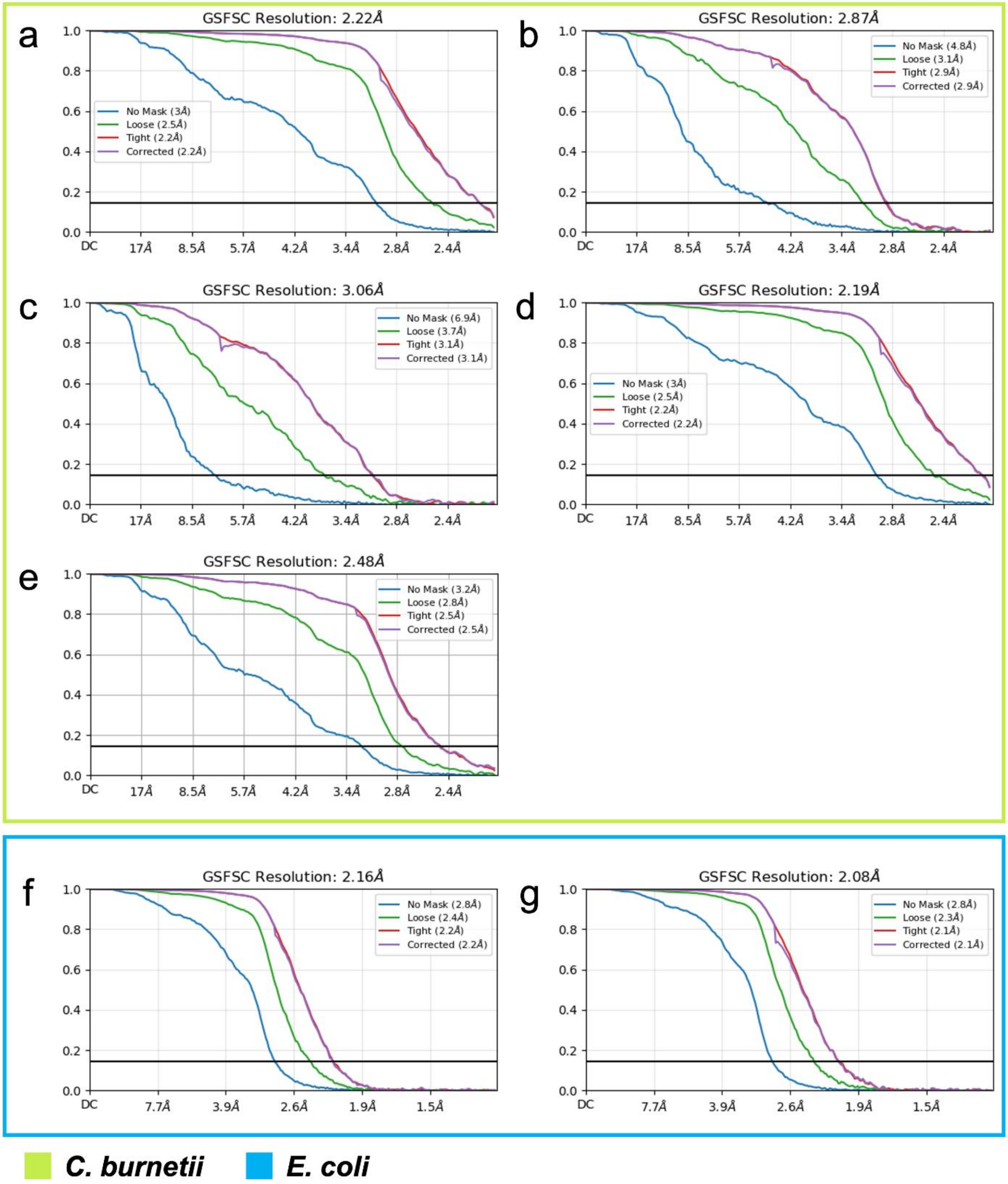

***Supplementary discussion***

#### $HPF_{cold}$ is a new class of ribosome hibernation factor

HPF represents the first structure of a hibernation factor with a cold shock domain, representing a new class of ribosome hibernation factor family [12]. As such, we propose to name this class $HPF_{cold}$. It is structurally distinct from existing HPF family members; $HPF_{long}$, $HPF_{short}$ or ribosome associated inhibitor A (RaiA) (Figure S2c). These are the primary hibernation factors in bacteria and dictate the specific form of ribosome hibernation that occurs. In gram-negatives, 100S ribosome dimers are formed through a combination of $HPF_{short}$ and ribosome modulation factor (RMF). $HPF_{short}$ occupies the tRNA binding cleft with RMF driving dimerising through a copy on the partner ribosome [4]. In other species, both roles are carried out by $HPF_{long}$, dimerising through its C-terminal domain (CTD) via an elongated linker [5]. 70S hibernation occurs through RaiA, with a C-terminal extension preventing 100S formation by clashing with RMF [4]. *C. burnetii* lacks a gene for both $HPF_{long}$ and RMF, with its only other known hibernation factor (*CBU_0745*) annotated as RaiA. However, *CBU_0745* lacks the characteristic C-terminal extension of RaiA, instead appearing to classify as $HPF_{short}$. The prokaryotic 100S dimerization interface falls upon the mRNA entrance tunnel region, conferring protection not provided by $HPF_{short}$. Loss of 100S hibernation is therefore associated with lower virulence and survival rates, with increased susceptibility to RNAses [13]. RMF confers protection to the 3' r16S [14]; it appears that $HPF_{cold}$ provides similar protection since $HPF_{cold}$ CSD binds and stabilises the anti-Shine Dalgarno sequence, blocking RNAse access. $HPF_{cold}$ may confer a high enough degree of protection from RNAses to keep ribosomes viable over a long time period, presumably a crucial capability for *C.*

*burnetii* during the extended hibernation periods required of the SCV form [15]. We were unable to identify ribosome dimers in our single particle data (although it is feasible any dimerised ribosomes would be lost during sample preparation). While an as yet unidentified factor (or even $HPF_{cold}$ CSD dimerization) could induce 100S assemblies, $HPF_{cold}$ may render this need obsolete by providing 100S-like capabilities in a 70S form. A transposon mutagenesis screen suggests $HPF_{cold}$ is not essential for *C. burnetii* growth in media at 37 °C [16]. Future experiments investigating the effects of a $HPF_{cold}$ knockout would be valuable for delineating its role in the bacterial hibernation and stress response, in *C. burnetii* and other prokaryotes.

Across the phylogenetic tree (Fig. 2f), the PF02482 (sigma-54) and PF00313 (cold shock domain) fusion is present in all major proteobacterial classes but with uneven distribution, consistent with either lineage-specific expansions or sampling biases. Fused-domain homologs are also present in Cyanobacteria and several non-proteobacterial phyla, producing a polyphyletic distribution. Mapping metabolic annotations onto the tree shows the fusion recurring in specialist guilds such as nitrifiers, anammox bacteria and multiple phototrophic lineages suggesting repeated recruitment into redox-intensive lifestyles across oxic–anoxic gradients and, therefore, ecological niche expansion rather than a single constrained function. Taken together, this pattern is consistent with a mix of processes: repeated, independent PF02482–PF00313 fusion events, horizontal transfers of an existing fusion, and lineage-specific losses, with retention where the fused domain architecture is advantageous. Within Gammaproteobacteria, *Coxiella* sequences form a distinct, well-supported subclade (Coxiellaceae/Legionellales), clearly separated from other gammaproteobacterial homologs. This placement points to lineage-specific functional constraints under an intracellular lifestyle rather than a recent cross-phylum acquisition. Notably, Alphaproteobacteria display the broadest radiation, with multiple sub-clusters that could reflect diversification of the fusion within this class or simply denser genome sampling. Conversely, the absence of the fused architecture in several major bacterial groups in our dataset points to niche-specific utility and possible functional constraints on maintaining this two-domain fusion. Together, these patterns place $HPF_{cold}$ in a clear phylogenetic context and support a scenario where the fusion has a long evolutionary history punctuated by clade-specific expansions and losses.

##### A doxycycline triple stack blocks the *C. burnetii* ribosome exit tunnel

The *C. burnetii* ribosomal exit tunnel in the 50S subunit is filled by a stack of three doxycycline molecules (Fig. 4a-c). This stack relies upon a magnesium induced rotated amide conformation (Fig. S10a), as has been observed in certain small molecule crystallisation conditions [17]. In comparison to the unbound structure, DOX2 stabilises r23S RNA U2506 (U2506) (Fig. S9c-d), involved in sensing ribosomal stalling [18]. U2506 takes a 'luminal', stalled-like conformation showing a rotation which has broken the hydrogen bond pair to G2603 (G2583). Additionally, A2082 (A2062), which is vital for sensing translational arrest [19] is stabilised by DOX1 through a Pi stack of the amide group (Fig. S9e, S10a).

Sarecycline bound to the *C. acnes* ribosome most closely resembles the DOX1 binding mode, however the *C. acnes* phosphate backbone conformation between 2629-2630 (2609-2610) would abrogate binding of doxycycline molecule 2 (Fig. S9c-d). Sarecycline is also too distant to interact with U2506 [8]. Like DOX1, tetracenomycin X stacks on U2606 but is instead rotated, moving out of range of the magnesium ion DOX1 utilises (Fig. S6e) [9]. Interestingly, some elements of macrolide binding are re-capitulated by the doxycycline stack, notably, the single conserved hydrogen bond required for macrolide binding [10]. In macrolides, the desosamine 2-OH group hydrogen bonds to A2078 (A2058). This role is taken by a water coordinated by the DOX2-3 magnesium, H-bonding to the N1 of A2078 (A2058) (Fig. S6f-g). Dimethylation of *E. coli* A2058 confers macrolide resistance through the disruption of a water mediated interaction. This same interaction is not imitated in the doxycycline binding, which may enable doxycycline to bind even in the presence of a dimethylated A2078 [10].

Erythromycin resistance is conferred by the ErmCL peptide, which interacts with U2506 to trigger translational arrest [18]. The tunnel blockage by doxycycline may preclude a similar peptide-based resistance mechanism. Whilst resistance to many antibiotics targeting the PTC (chloramphenicol, linezolid etc) is conferred by methylation of 2503 C8 [20, 21], given the distance from the bound doxycycline molecules, this would be unlikely to obstruct *C. burnetii* doxycycline binding (Fig. S8e). We also note that *C. burnetii* appears to lack a gene encoding a tetracycline destructase, a recently identified family of bacterial resistance enzymes [22, 23].

We also note that while our results in *C. burnetii* were observed with three doxycycline molecules, we see no reason to constrain this to a single antibiotic. In the first case, combinatorial therapy with existing, licensed tetracycline family members may further optimise the binding within the stack. This could concomitantly achieve higher affinities and most crucially, would sidestep extensive development and clinical trial processes. It may be that different combinations would be optimal for different organisms. Subsequent structure guided development of new molecules, fully exploiting each binding site within the stack could yield a species-specific gold standard, most effective treatment.

**Cryo-EM reveals multiple mechanisms of ribosome inhibition by doxycycline**

**Stuart et al.**

#### *References*


1. Abramson, J., et al., *Accurate structure prediction of biomolecular interactions with AlphaFold 3.* Nature, 2024. **630**(8016): p. 493-500.
2. Schindelin, H., et al., *Crystal structure of CspA, the major cold shock protein of Escherichia coli.* Proc Natl Acad Sci U S A, 1994. **91**(11): p. 5119-23.
3. Pettersen, E.F., et al., *UCSF ChimeraX: Structure visualization for researchers, educators, and developers.* Protein Sci, 2021. **30**(1): p. 70-82.
4. Polikanov, Y.S., G.M. Blaha, and T.A. Steitz, *How hibernation factors RMF, HPF, and YfiA turn off protein synthesis.* Science, 2012. **336**(6083): p. 915-8.
5. Franken, L.E., et al., *A general mechanism of ribosome dimerization revealed by single-particle cryo-electron microscopy.* Nat Commun, 2017. **8**(1): p. 722.
6. Sievers, F., et al., *Fast, scalable generation of high-quality protein multiple sequence alignments using Clustal Omega.* Mol Syst Biol, 2011. **7**: p. 539.
7. Robert, X. and P. Gouet, *Deciphering key features in protein structures with the new ENDscript server.* Nucleic Acids Res, 2014. **42**(Web Server issue): p. W320-4.
8. Lomakin, I.B., et al., *Sarecycline inhibits protein translation in Cutibacterium acnes 70S ribosome using a two-site mechanism.* Nucleic Acids Res, 2023. **51**(6): p. 2915-2930.
9. Osterman, I.A., et al., *Tetracenomycin X inhibits translation by binding within the ribosomal exit tunnel.* Nat Chem Biol, 2020. **16**(10): p. 1071-1077.
10. Svetlov, M.S., et al., *Structure of Erm-modified 70S ribosome reveals the mechanism of macrolide resistance.* Nat Chem Biol, 2021. **17**(4): p. 412-420.
11. Punjani, A., et al., *cryoSPARC: algorithms for rapid unsupervised cryo-EM structure determination.* Nat Methods, 2017. **14**(3): p. 290-296.
12. Prossliner, T., et al., *Ribosome Hibernation.* Annu Rev Genet, 2018. **52**: p. 321-348.
13. Gohara, D.W. and M.F. Yap, *Survival of the drowsiest: the hibernating 100S ribosome in bacterial stress management.* Curr Genet, 2018. **64**(4): p. 753-760.
14. Prossliner, T., et al., *Hibernation factors directly block ribonucleases from entering the ribosome in response to starvation.* Nucleic Acids Res, 2021. **49**(4): p. 2226-2239.
15. Kersh, G.J., et al., *Presence and persistence of Coxiella burnetii in the environments of goat farms associated with a Q fever outbreak.* Appl Environ Microbiol, 2013. **79**(5): p. 1697-703.
16. Metters, G., et al., *Identification of essential genes in Coxiella burnetii.* Microb Genom, 2023. **9**(2): p. 944.
17. Santos, O.M., Silva, D.M., Martins, F.T., Legendre, A.O., Azarias, L.C., Rosa, I.M., Neves, P.P., de Araujo, M.B. and Doriguetto, A.C., *Protonation pattern, tautomerism, conformerism, and physicochemical analysis in new crystal forms of the antibiotic doxycycline.* Crystal growth & design, 2014. **14**(8): p. 3711-3726.
18. Koch, M., et al., *Critical 23S rRNA interactions for macrolide-dependent ribosome stalling on the ErmCL nascent peptide chain.* Nucleic Acids Res, 2017. **45**(11): p. 6717-6728.

19. Vazquez-Laslop, N., et al., *The key function of a conserved and modified rRNA residue in the ribosomal response to the nascent peptide.* EMBO J, 2010. **29**(18): p. 3108-17.
20. Kehrenberg, C., et al., *A new mechanism for chloramphenicol, florfenicol and clindamycin resistance: methylation of 23S ribosomal RNA at A2503.* Mol Microbiol, 2005. **57**(4): p. 1064-73.
21. Aleksandrova, E.V., et al., *Structural basis of Cfr-mediated antimicrobial resistance and mechanisms to evade it.* Nat Chem Biol, 2024. **20**(7): p. 867-876.
22. Blake, K.S., et al., *Sequence-structure-function characterization of the emerging tetracycline destructase family of antibiotic resistance enzymes.* Commun Biol, 2024. **7**(1): p. 336.
23. Forsberg, K.J., et al., *The Tetracycline Destructases: A Novel Family of Tetracycline-Inactivating Enzymes.* Chem Biol, 2015. **22**(7): p. 888-97.
